# Targeted suppression of type 1 interferon signaling during RNA delivery enhances vaccine-elicited immunity

**DOI:** 10.1101/2025.03.09.642244

**Authors:** B.J. Kim, Ryan R. Hosn, Tanaka K. Remba, Jonathan Dye, Howard H. Mak, Jae Yun Jeong, Milton Cornwall-Brady, Wuhbet Abraham, Laura Maiorino, Mariane B. Melo, Bridget Li, Yuebao Zhang, Yizhou Dong, Darrell J. Irvine

**Author notes:** Department of Immunology & Microbiology, The Scripps Research Institute, La Jolla, CA, 92037, USA (Current address).

## Abstract

RNA vaccines have emerged as a breakthrough technology, and one promising modality employs alphavirus-derived self-replicating RNA (repRNA) to express vaccine antigens. However, both the lipid nanoparticles (LNP) commonly used to deliver RNA and virus-like amplification of repRNAs trigger innate immune recognition, especially via type I interferon (IFN) signaling. To modulate IFN responses during vaccination, we formulated LNPs co-delivering antigen-encoding RNA together with siRNA targeting the interferon-α/β receptor-1 (IFNAR1). siRNA-mediated repression of IFNAR1 increased antigen expression from repRNAs by >10-fold, increased immune cell infiltration, and increased antigen presenting cell activation in the injection site and draining lymph nodes. Compared to repRNA alone, siRNA/repRNA co-delivery increased serum antibody titers >10-fold, dramatically augmented antigen-specific germinal center (GC) B cell responses, and primed 4.4-fold more antigen-specific T cells. *Ifnar1* silencing by siRNA co-delivery similarly enhanced mRNA vaccines. Thus, siRNA co-delivery is a readily translatable approach to substantially enhance the immunogenicity of RNA vaccines.

## MAIN TEXT

Effective delivery of antigen-encoding mRNA in humans using lipid nanoparticles (LNPs) represents a significant advancement in vaccinology.^1,2^ The development of self-replicating RNA (replicon/repRNA) technology has the potential to further increase the potency of RNA vaccines and reduce the required RNA (and LNP) dosage, thereby also enhancing vaccine safety.^3–10^ RepRNAs can provide very high levels of payload expression in transfected cells, as they encode both a subgenomic payload gene of interest (e.g., an antigen) and non-structural proteins that copy both the complete RNA strand and the subgenome, leading to self-amplification. Countering this process are multiple pathways of innate immune recognition: repRNA may be recognized by toll-like receptor-7 (TLR7) following LNP uptake into endosomes, while replicons released into the cytoplasm can be recognized by RIG-I and MDA-5.^6,11–15^ In addition to the RNA itself, LNPs commonly used to deliver both mRNA and repRNA vaccines themselves trigger NF-κB activation^16,17^ and type 1 interferon (IFN) responses^18–20^. Mechanistic insights into how LNPs cause such responses are under active research, such as LNP-mediated endosomal rupture^21^ and inflammasome activation by ionizable lipids^22^. These pathways for innate immune recognition can hinder the ultimate vaccine-elicited response by inhibiting RNA replication and translation in anti-viral defense^15,23–25^, but may also have an adjuvanting role, stimulating the production of inflammatory cytokines and recruitment of immune cells to the vaccination site.^26,27^

A key output of these RNA recognition pathways is the production of type 1 IFNs, which can act in both an autocrine and paracrine manner.^28,29^ Type 1 IFN production has been shown in a number of studies to be a key regulator of repRNA expression *in vivo*: intramuscular immunization of mice with repRNA in IFN-α/β receptor 1 knockout (*ifnar*^-/-^) animals led to 100-fold increases in reporter gene expression compared to WT animals.^11^ Others have also reported a 10-fold increase in luciferase expression in *ifnar*^-/-^ mice.^30^ However, the effect of global IFN signaling deficiency on the immunogenicity of replicon vaccines is less clear: immunization of *ifnar*^-/-^ vs. WT animals with repRNA encoding various viral antigens has been reported to elicit either no statistically significant changes in antibody/neutralizing titers^11,31,32^, or substantial increases in antibody responses^30^. Some studies reported no changes in CD4^+^ T cell responses^11,30^, and modest increases in CD8^+^ T cell responses^11,30,31^. These highly varied responses are similar to effects observed with mRNA, where both increases^24,25^ and decreases^33,34^ in T cell responses have been reported in IFN-deficient animals. Interestingly, type 1 IFN signaling was found to be critical for the activation of CD8^+^ T cells^34^, but was not required for dendritic cell recruitment in response to RNA vaccines^25^. It seems likely that the varied/limited effects seen in these studies reflect the complex roles of type I interferons in the immune response, which extend well beyond the RNA-transfected cell.^24^

Here, we developed an approach to focus modulation of type I IFN signaling on transfected cells *in vivo*, by co-delivering siRNA targeting *Ifnar1* together with repRNA or modRNA in the same LNP. Using RNA vaccines encoding a clinically-relevant HIV immunogen– a transmembrane form of a germline targeting HIV Env SOSIP trimer– we assessed the impact of *in vivo Ifnar1* silencing on repRNA expression, changes in the composition of immune cells recruited at the injection site and draining lymph nodes, antigen presenting cell (APC) activation, and downstream cellular and humoral immune responses. These studies revealed that co-delivery of RNA vaccines with siRNA against *Ifnar1* enhances antigen expression, immune cell infiltration and stimulation, and diverse downstream antigen-specific vaccine responses.

## RESULTS

### Designing RNA vaccines to query the role of IFNAR1 in vaccine-elicited immunity

As a model system, we synthesized repRNA encoding a transmembrane form of a germline targeting HIV Env trimer termed N332-GT2.^35^ This Env trimer immunogen is designed to prime B cells in humans capable of evolving to produce broadly neutralizing antibodies targeting the V2-Apex neutralizing site on HIV; a closely related trimer is currently in phase I human testing.^36^ To determine whether type I IFN signaling impacts the development of antigen-specific B cells in germinal centers (GCs) following RNA immunization, we vaccinated wildtype and *ifnar*^-/-^ mice with LNPs carrying repRNA encoding N332-GT2. Two weeks post-vaccination, we observed that while there were no significant differences in the total frequency of germinal center (GC) B cell or follicular helper T (T_fh_) cell populations (**Extended Data Fig. 1a-b**), antigen-specific GC B cells that were present at a frequency of less than 1% in WT animals were increased 9.4-fold in *ifnar*^-/-^ mice (**Extended Data Fig. 1c-d**). Further, 60% of these animals had also already seroconverted, while antibody titers in WT animals were still at baseline at this timepoint (**Extended Data Fig. 1e-f**). Thus, the type 1 IFN pathway appears to play an important role in restraining humoral responses to repRNA.

We hypothesized that co-delivery of repRNA together with siRNA targeting IFN sensing might provide a means to pharmacologically modulate the interferon response in a manner focused on RNA-transfected cells without global IFN suppression, and over a time window that would be relevant for amplifying repRNA vaccine responses. To test this idea, we prepared (1) LNPs carrying repRNA alone, (2) LNPs co-loaded with repRNA and one of two different siRNAs targeting *Ifnar1* or a control scrambled siRNA sequence (siScramble), and (3) a “cocktail” case consisting of LNPs carrying repRNA alone mixed with LNPs carrying siRNA (**Fig. 1a**). LNPs were synthesized using microfluidic mixing of lipids in ethanol with aqueous phase repRNA in citrate. For these initial studies, we utilized an LNP formulation containing two ionizable lipids, TT3^37^ and DLin-MC3-MDA^38^, which provided more efficient transfection of mouse muscles with repRNA compared to LNP compositions employed in the Moderna (MDR) or Pfizer (PFZ) COVID-19 mRNA vaccines (**Supplementary Table S1**, **Extended Data Fig. 2a-b**).

**Figure 1.**
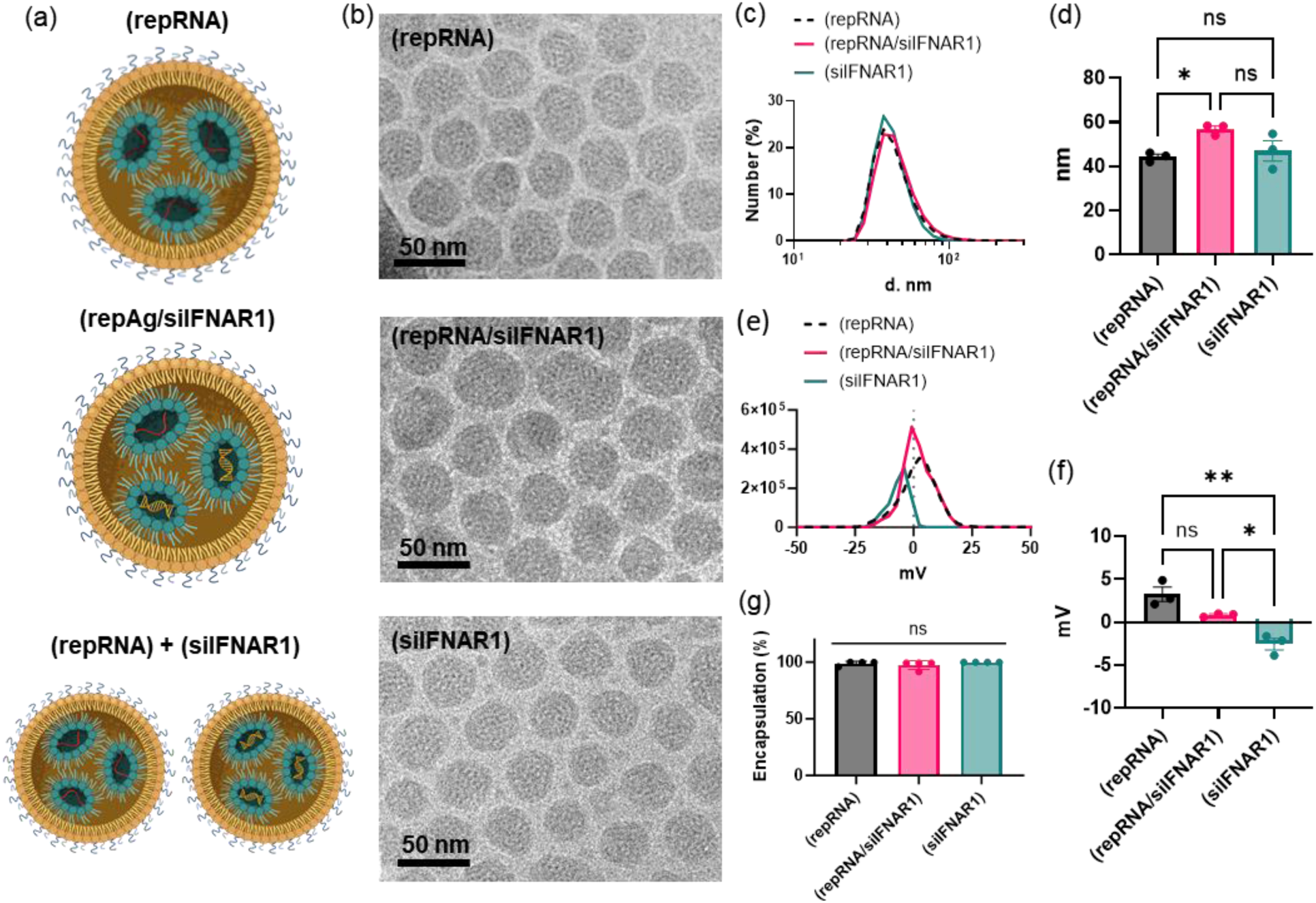
Formulation of LNPs for co-delivery of repRNA and siRNA. (a) Schematic of vaccine groups: ‘(repRNA)’ shows an LNP loaded with replicons (red strands), ‘(repRNA/siIFNAR1)’ shows an LNP that co-loads replicons and siRNA against *Ifnar1* (yellow double helices), and ‘(repRNA) + (siIFNAR1)’ shows a mixture of LNPs that are loaded with either replicons or siRNA against *Ifnar1*; (b) Cryogenic electron microscope (Cryo-EM) images of: (repRNA), (repRNA/siIFNAR1), and (siIFNAR1); (c) Dynamic light scattering (DLS)-measured number-weighted size distribution of the three LNP vaccines; (d) Average hydrodynamic diameter of the three LNP vaccines; (e) Zeta-potential measurements of the three LNP vaccines; (f) Average zeta-potential of the three LNP vaccines; and (g) RNA encapsulation efficiency of the three LNP vaccines. Error bars represent standard error means (s.e.m.) and statistics represent one-way ANOVA and Tukey’s HSD Test (ns= not significant, * *p<*0.05; **, *p*<0.01).

We characterized LNPs loaded with repRNA alone, repRNA together with an siRNA targeting *Ifnar1* (siIFNAR1), or LNPs loaded with only siIFNAR1 (**Supplementary Table S2**). For the co-loaded formulation, we selected a repRNA:siRNA mass ratio of 1:2 based on enhanced early replicon expression observed when delivering repRNA expressing luciferase as a reporter gene (**Extended Data Fig. 2c**). Cryo-EM and dynamic light scattering showed that all three formulations formed relatively monodisperse populations of nanoparticles approximately 40 nm in diameter, with near-neutral zeta potentials and high RNA encapsulation efficiencies (**Fig. 1b-g**).

### Silencing *Ifnar1* augments repRNA expression *in vitro*

C2C12 mouse myoblast cells transfected with LNPs carrying GFP-encoding repRNA alone, repRNA with siRNA, or mixtures of LNPs carrying only repRNA and only siRNA all showed similar high levels of cell viability and LNP uptake (**Fig. 2a-b**). We compared knockdown of *Ifnar1* using two different co-delivered siRNA sequences (siIFNAR1.1 and siIRNFAR1.2), and characterized replication of the RNA by qRT-PCR using primers targeting the *nsP3* domain of the backbone and the subgenomic *Gfp* payload (**Fig. 2c**, **Supplementary Table 3**). LNPs co-loaded with replicon and siIFNAR1 triggered over 95% knockdown of *Ifnar1*, while LNPs carrying repRNA and control scrambled siRNA led to no significant change, and co-delivery of separate LNPs carrying repRNA and siRNA showed *Ifnar* knockdown but with lower efficiency (77%, **Fig. 2d**). Knockdown of *Ifnar1* in cells receiving siRNA/repRNA co-delivery was accompanied by a significant increase in copies of the repRNA backbone (over 6-fold increase in *nsP3* transcripts for siIRNAR1.2, **Fig. 2e**) and up to 17-fold increases in copies of the replicon payload *Gfp* (**Fig. 2f**) compared to cells transfected with replicon alone or co-delivered with siScramble. At the protein level, the mean fluorescence intensity (MFI) of GFP-expressing cells also significantly increased when cells were transfected with LNPs co-delivering repGFP and siIFNAR1, up to 1.7-fold over repRNA alone for the optimal siIFNAR1.2 sequence (**Fig. 2g-h**). These results indicate that replicon/siRNA co-delivery can be used to substantially enhance both repRNA genome replication as well as copies of payload gene transcripts without compromising cell viability.

**Figure 2.**
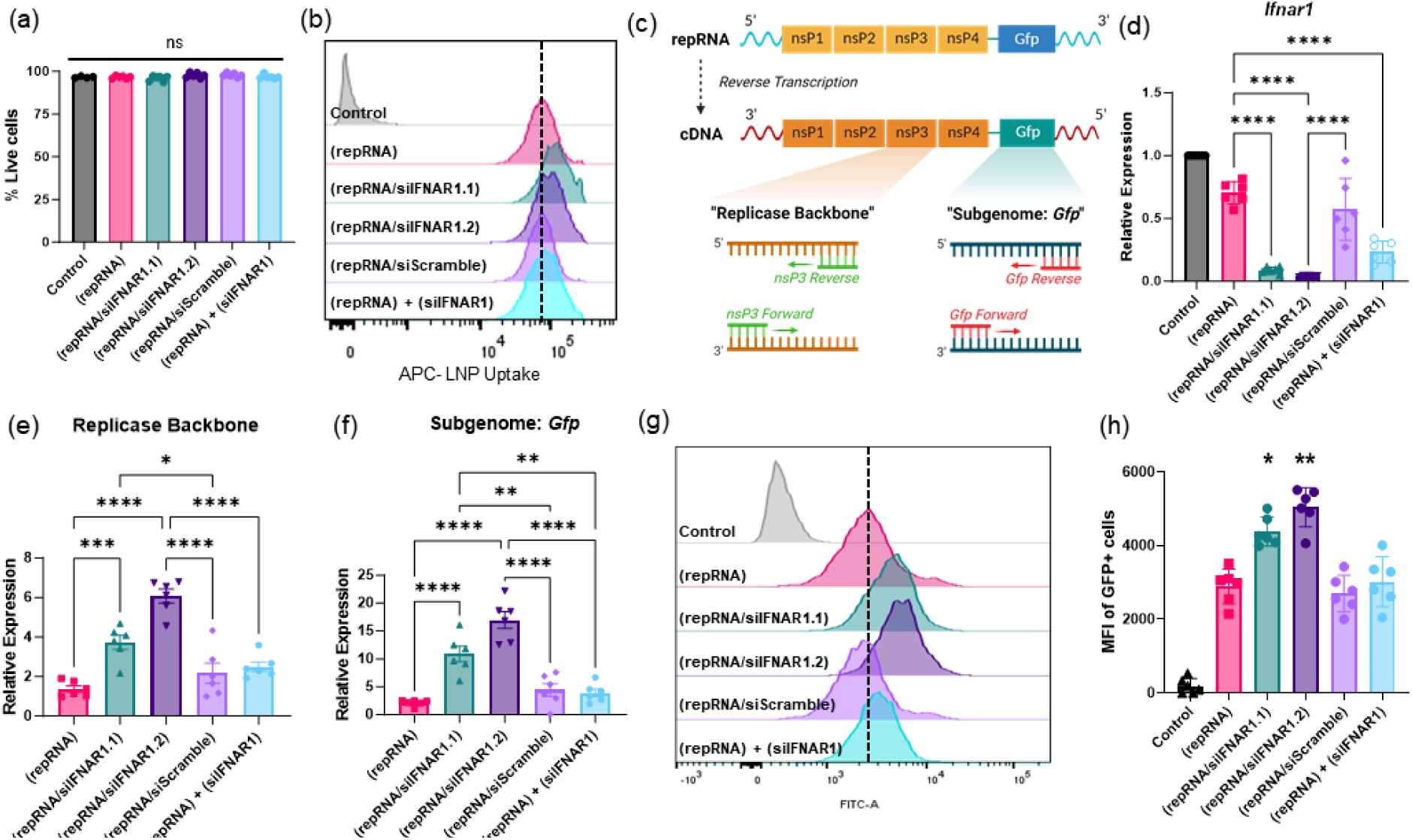
Silencing *Ifnar1* in myoblasts enhances expression of repRNA *in vitro*. C2C12 mouse myoblasts were treated with PBS, LNPs loaded with repRNA encoding for GFP (repGFP), LNPs that co-load repGFP and siRNA against *Ifnar1* (repRNA/siIFNAR1.1 or repRNA/siIFNAR1.2), LNPs that co-load repGFP and control scrambled siRNA (repRNA/siScramble), or a mixture of LNPs loaded with only repRNA and LNPs loaded with only siIFNAR1.2 (repRNA) + (siIFNAR1). (a-b) Transfected C2C12 mouse myoblasts were analyzed by flow cytometry at 24h. Shown are cell viability (a) and uptake of fluorescently-labeled LNPs as representative histograms (b). (c) Schematic illustrating the replicon gene makeup, and binding sites for primers used in (e) and (f). (d-f) At 24h post-transfection, cells were lysed for qRT-PCR analysis of RNA expression. Shown are *Ifnar1* (d); repRNA backbone (e); and *Gfp* (f). (g-h) Flow cytometry analysis of transfected cells at 24h, showing representative histograms of GFP expression (g); and MFIs of GFP-expressing live cells (h). Samples are presented as means ± standard error means (s.e.m.). Statistics represent one-way ANOVA and Tukey’s HSD Test (ns= not significant; *, p<0.05; **, *p*<0.01; ***, *p*<0.001; ****, *p<*0.0001).

### siRNA/repRNA co-delivery enhances replicon expression *in vivo*

Next, we evaluated effects of delivering *Ifnar1*-targeting siRNA on replicon expression *in vivo*. Balb/c mice were vaccinated intramuscularly (i.m.) in the gastrocnemius with LNPs carrying repRNA encoding GFP (“(repRNA)”), siIFNAR1.2 co-loaded LNPs (“(repRNA/siIFNAR1)”), or a mixture of the repRNA-loaded and siIFNAR1.2-loaded LNPs (“(repRNA) + (siIFNAR1)”). qRT-PCR analysis of injected muscles showed that while administration of replicon alone led to a 6.7-fold increase in *Ifnar1* expression 1-day post-administration, this upregulation was completely blocked for at least 3 days following replicon/siRNA co-delivery (**Fig. 3a**). Interestingly, *Ifnar1* expression then rebounded at day 7 significantly to ∼3-fold higher expression than replicon-only treatment, which had dropped substantially by this timepoint. This suppression of *Ifnar1* correlated with >10-fold increased transcripts for the repRNA backbone and its subgenome payload, which were maintained over time (**Fig. 3b-c**). Although co-treatment with repRNA and siRNA delivered by separate LNPs also blocked *Ifnar1* upregulation at early times, this treatment failed to elicit the same increases in replicon backbone and *Gfp* transcript levels (**Fig. 3a-c**). In draining popliteal lymph nodes (dLNs), *Ifnar1* expression was transiently upregulated at day 1 post-RNA administration (**Fig. 3d**); siRNA co-delivery reduced this upregulation compared to replicon alone by 2-fold, accompanied by 4.4-fold and 5.5-fold increased peak levels of the replicon backbone and subgenome transcripts, respectively, in the dLN at day 3 (**Fig. 3e-f**).

**Figure 3.**
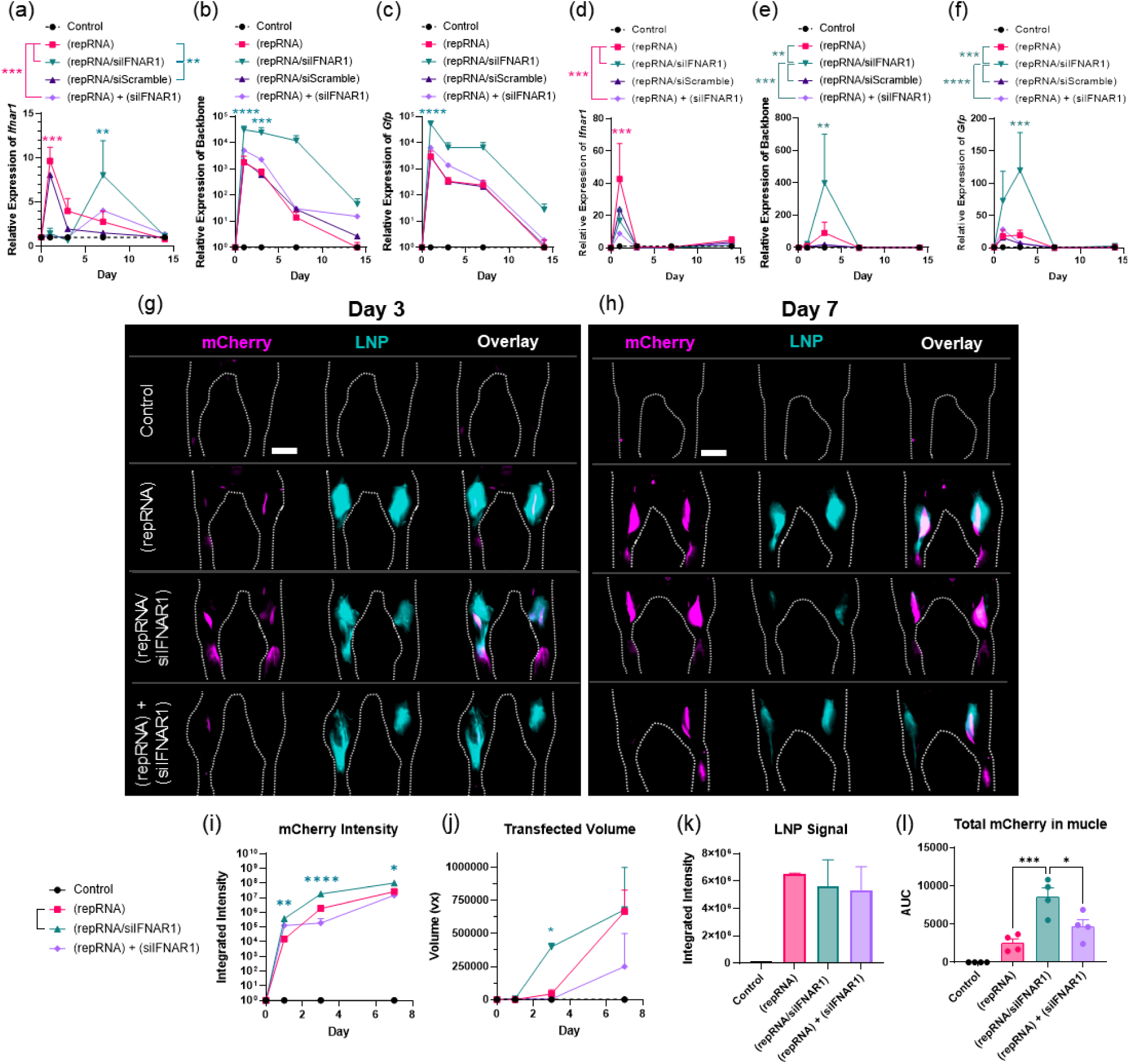
*In vivo* knockdown of *Ifnar1* increases expression level of repRNA when packaged and delivered with siIFNAR1 in the same LNP. Groups of Balb/C mice (n=5 animals/group) were immunized i.m. in each leg with 1 µg repRNA loaded in LNPs, and sacrificed at days 1, 3, 7, and 14 post-injection for gastrocnemius (muscle) and popliteal lymph node (pLN) harvesting. RNA was purified from the tissues for qRT-PCR-based quantification of the relative expression levels in the muscle of: (a) *Ifnar1*; (b) the repRNA backbone encoding for the replicase; and (c) *Gfp* as the model repRNA payload. From the pLN, we show (d) *Ifnar1*; (e) the repRNA backbone; and (f) *Gfp*. Bars represent standard error means. (g-m) Balb/C mice were administered i.m. in each leg with 1 µg mCherry-encoding repRNA loaded in DiD-labelled LNPs. Cryo-fluorescence tomography (CFT) was used to visualize the lower limbs of mice for expression of repRNA encoding for mCherry (magenta) and LNPs labelled with DiD (cyan) at: (g) day 3 post-i.m. injection; and (h) day 7 post-i.m. injection. Expression intensity of mCherry (i), as well as the volume of mCherry expression (j), and signal intensity of DiD-labeled LNPs (k) were quantified and plotted as a function of integrated intensity or volume (voxel; vx). (l) ELISA-quantification of mCherry expression as area under the curve of mCherry protein levels at day 7-post i.m. injection. Scale bars in (g) and (h) represent 5 mm. Data are shown as means ± s.e.m. Statistics represent one-or two-way ANOVA and Tukey’s HSD Test (ns= not significant; *, p<0.05; **, p<0.01; ***, p<0.001; ****, p<0.0001). Statistics in (j) and (l) represent significant difference over the (repRNA) group.

We next used cryofluorescence tomography (CFT)– where frozen tissues are serially sectioned and imaged using fluorescence microscopy, followed by digital reconstruction to enable whole-tissue 3D fluorescence imaging– to visualize the muscle injection site following delivery of fluorescent LNPs carrying replicons encoding the reporter gene mCherry. As shown in 2D maximum-intensity projections of the 3D imaging data (**Fig. 3g-h**) and in volumetric quantification of mCherry signal (**Fig. 3i-j**), replicon/siRNA co-administration led to a 10-fold higher integrated intensity of mCherry expression (**Fig. 3i**) in an 11-fold greater volume of the muscle tissue (**Fig. 3j**) at day 3 compared to immunization with repRNA alone. Reporter gene expression increased over time for repRNA alone, but repRNA/siRNA co-delivery maintained a 3.8-fold higher integrated intensity of mCherry over the repRNA-alone group 7 days post-vaccination. Notably, delivery of repRNA LNPs mixed with siRNA LNPs led to lower mCherry expression signal and transfected volume compared to delivery of the replicon and siRNA together in the same LNP (**Fig. 3g-j**). As expected, the total LNP signal in the tissues was not statistically different between groups (**Fig. 3k**). We next quantified mCherry expression in the muscles over time by ELISA analysis of tissue lysates. Similar to the findings from CFT, we saw found 3.4-fold higher levels of mCherry expressed for repRNA/siIFNAR1 co-loaded LNPs compared to repRNA alone at day 7 post-administration (**Fig. 3l**). Altogether, silencing of the type I IFN responsiveness through *Ifnar1* knockdown substantially amplified expression of replicon-encoded payload genes over at least the first week post administration.

### Suppression of *Ifnar1* enhances immune cell recruitment and stimulation in muscle and draining lymph nodes

We next characterized the impact of *Ifnar1* silencing on immune cell recruitment/phenotypes in the muscle and dLNs by flow cytometry (**Supplementary Fig. S1**). As LNPs induce a variety of cytokines, we expected that *Ifnar1* silencing would not block cytokine/chemokine expression in injected muscles. This was confirmed by multiplex ELISA analysis of muscle lysates following repRNA administration, where we saw that IL-1β, TNF-α, and IL-6 were all induced for both repRNA alone and repRNA/siRNA co-delivery (**Extended Data Fig. 3a-c**), and some chemokines were actually induced in the muscle by repRNA/siRNA co-delivery at *higher* levels than repRNA delivery alone (**Extended Data Fig. 3d-f**). Immune cell infiltration in the muscle was detected by 3 days post-vaccination, and interestingly, repRNA/siRNA co-delivery amplified immune cell recruitment, reaching 7-fold higher levels vs. repRNA alone at day 7 (**Fig. 4a**). By day 7, repRNA/siIFNAR1 co-delivery had increased the recruitment of cDC1 dendritic cells (∼8-fold, **Fig. 4b**), cDC2 dendritic cells (∼3-fold, **Fig. 4c**), and monocytes (∼13-fold, **Fig. 4d**), as well as neutrophils and macrophages (**Extended Data Fig. 4a-b**). Interestingly, adaptive immune cells were also enriched at the injected muscle (**Fig. 4e-f**), with B cells increased 20-fold over repRNA alone at day 7. By contrast, control immunization with repRNA/scrambled siRNA co-delivery or repRNA LNPs mixed with siRNA LNPs generally elicited immune cell infiltrates very similar to repRNA alone (**Extended Data Fig. 4c-h**). Among these recruited cells, mCherry expression was primarily detected in myeloid cells, and most prominently monocytes (**Fig. 4g**, **Extended Data Fig. 4i-k**). In addition to more transfected cells, repRNA/siRNA co-delivery vaccination yielded higher mean fluorescence intensities of mCherry expression on a per-cell basis (1.8-fold and 1.4-fold on days 3 and 7, respectively, **Fig. 4h**, **Extended Data Fig. 4k**).

**Figure 4.**
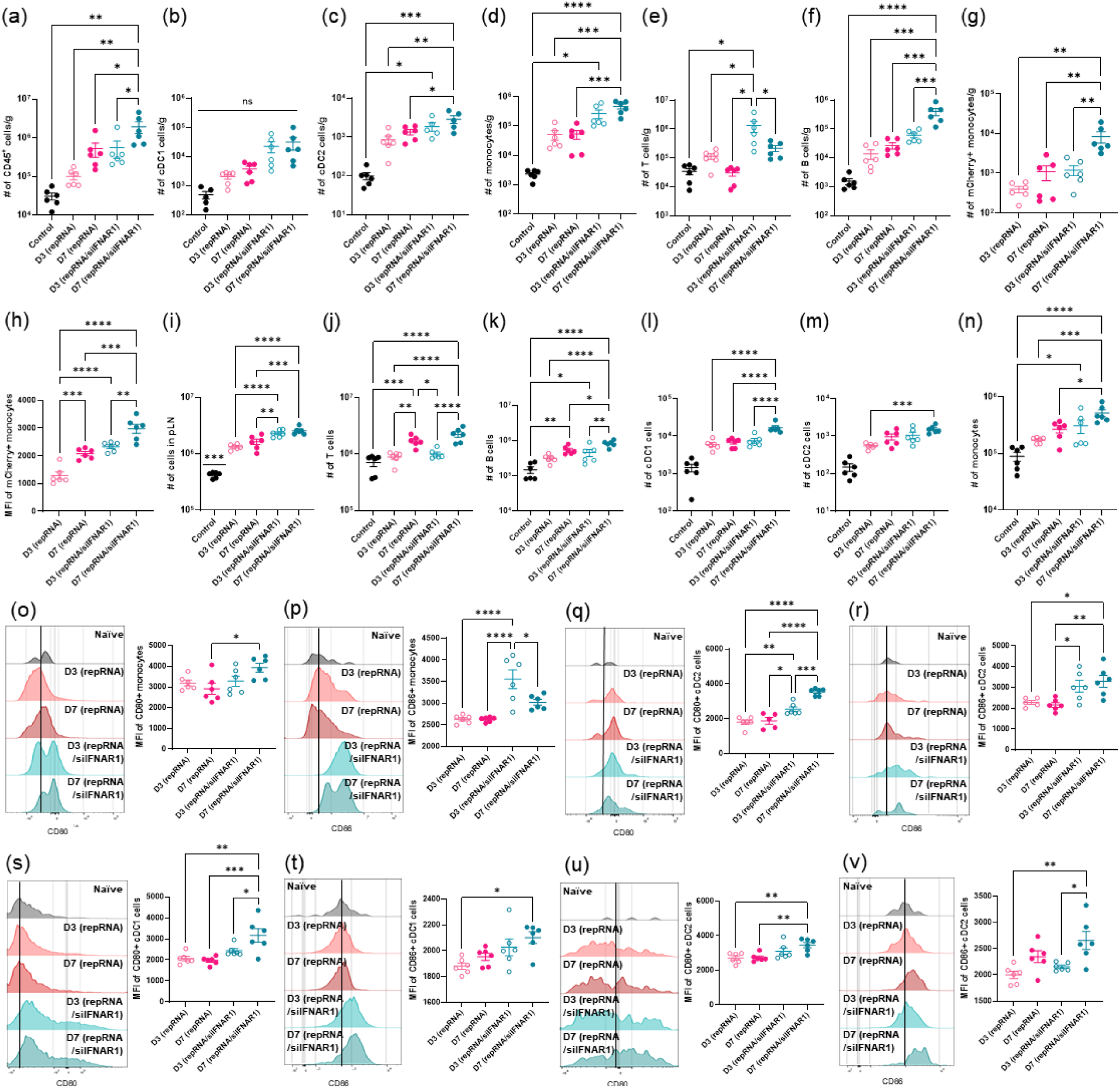
*Ifnar1* silencing increases immune cell infiltration and activation in the muscle and the draining lymph node. (a-d) Groups of Balb/C mice (n=5 animals/group) were immunized i.m. in each leg with 1 µg repRNA (encoding either immunogen or mCherry) loaded in LNPs, and sacrificed at days 3 and 7 post-injection for gastrocnemius muscle and popliteal lymph node harvesting. Single cells from muscles (a-h) and popliteal lymph nodes (i-n) were analyzed for flow cytometry-based immunophenotyping. Shown are counts from the muscle of: (a) CD45+ cells; (b) cDC1; (c) cDC2; (d) monocytes; (e) T cells; (f) B cells; (g) mCherry+ monocytes; and (h) MFI of mCherry+ monocytes. From the popliteal lymph node, we show counts of: (i) total cellularity; (j) T cells; (k) B cells; (l) cDC1s; (m) cDC2; and (n) monocytes. Cells were also assessed for activation by CD80 (o, q, s, u) and CD86 (p, r, t, v) markers. Shown are histograms and bar plot by MFI of the following cells from the muscle: (o) CD80 activation by monocytes; (p) CD86 activation by monocytes; (q) CD80 activation by cDC2s; and (r) CD86 activation by cDC2s. From the lymph node, we show: (s) CD80 activation by cDC1; (t) CD86 activation by cDC21; (u) CD80 activation by cDC2s; and (v) CD86 activation by cDC2s. Statistics represent one-way ANOVA and Tukey’s HSD Test (ns= not significant; *, p<0.05; **, p<0.01; ***, p<0.001; ****, p<0.0001).

Replicon vaccination induced expansion of draining lymph nodes, with *Ifnar1* silencing via siRNA co-delivery modestly increasing total cellularity vs. repRNA alone (∼1.6-1.8-fold higher, **Fig. 4i**). Among various cell subsets (**Fig. 4j-n**), siRNA co-delivery most impacted cDC1s, cDC2s, and monocytes, increasing each of these populations by ∼2-fold by day 7 (**Fig. 4l-n**).

Next, we evaluated activation of antigen presenting cells (APCs) in the muscle and dLN via expression of the key costimulatory receptors CD80 and CD86. Interestingly, immunization with replicon alone did not result in substantial increases in APC activation over naïve animals in either the muscle or dLN in the first week post-vaccination. However, vaccination with the replicon in conjunction with siRNA against *Ifnar1* yielded notable upregulation of CD80 and CD86 in monocytes (**Fig. 4o-p**) and cDC2s (**Fig. 4q**-**r**) in the muscle at day 7. In the draining lymph node, costimulatory receptor expression was significantly upregulated by day 7 in cDC1s (**Fig. 4s-t**), and cDC2s (**Fig. 4u-v**). Overall, we found that despite suppressing one inflammatory cue, co-delivery of replicon with siRNA against *Ifnar1* stimulated significantly higher infiltration of immune cells to the administration site and amplified APC activation in the muscle and dLNs.

### *Ifnar1* silencing enhances repRNA vaccine-elicited immune responses

We next assessed how these changes in gene expression and local inflammation relate to the immunogenicity of repRNA vaccines. We first immunized Balb/c mice with LNPs loaded with repRNA encoding N332-GT2 trimer with or without siIFNAR1.2, siScramble, or with the co-treatment vaccine. Flow cytometric analyses (**Supplementary Fig. S2a**) of cells recovered from the draining popliteal lymph nodes at day 14 revealed that vaccines co-delivering antigen-encoding repRNA and siIFNAR1 significantly increased expansion of total GC B cells and follicular helper T cells (Tfh, **Fig. 5a-b**). Most strikingly, staining with antigen tetramers revealed that siRNA co-delivery substantially amplified the antigen-specific GC B cell response, increasing both the total number of GC B cells capable of recognizing the intact antigen and the proportion of these cells that had class switched to IgG (**Fig. 5c-e**). Gating on GC B cells with similar IgG expression levels (**Supplementary Fig. S2b**), we found that antigen-specific B cells elicited by repRNA/siIFNAR1 co-delivery had an ∼8-fold higher mean fluorescence intensity of antigen binding (**Fig. 5f-g**), suggesting enhanced affinity maturation induced by *Ifnar1* silencing.^39^

**Figure 5.**
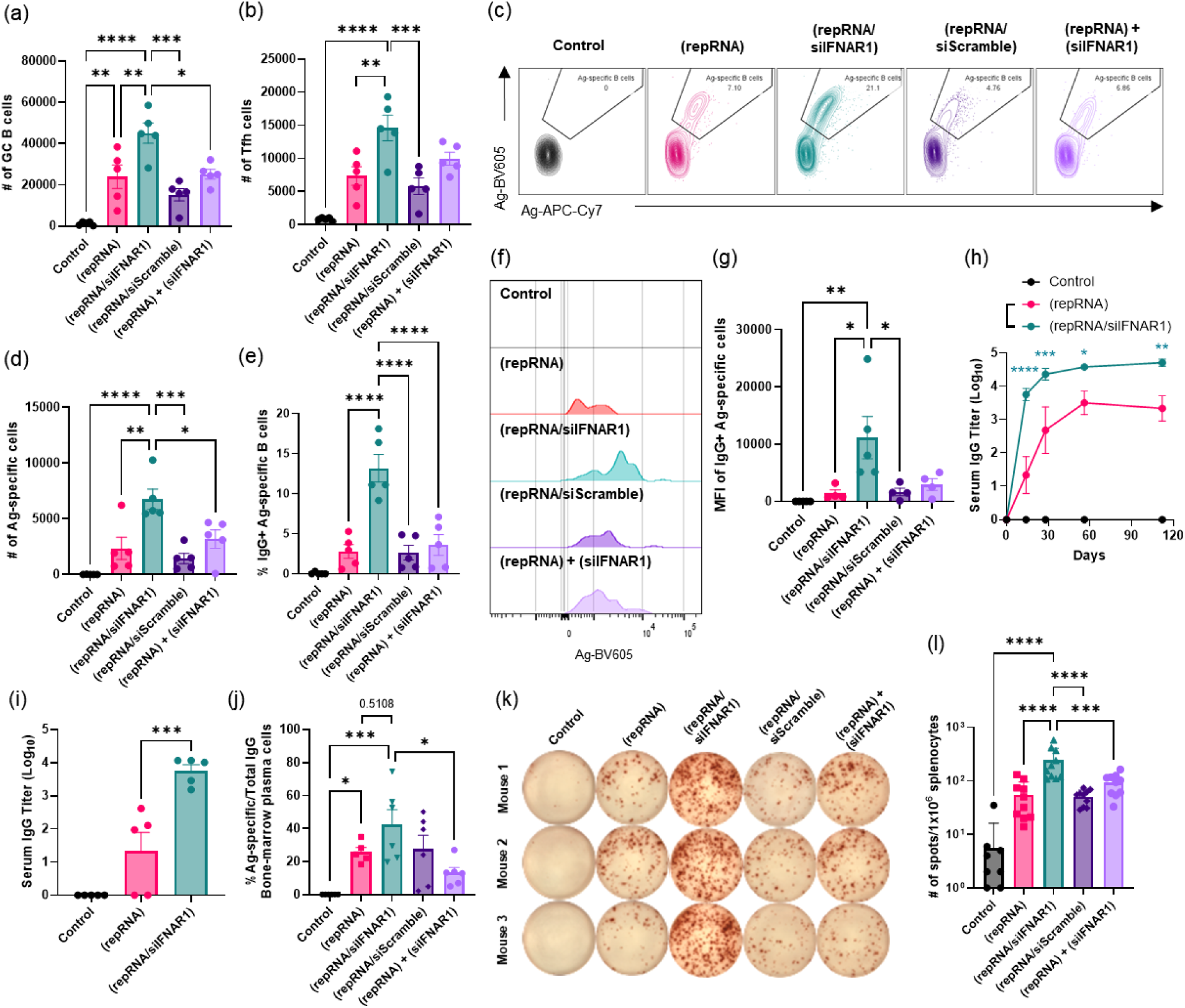
Silencing *Ifnar1* induces significantly more robust antibody production, antigen-specific B cell expansion, and antigen-specific T cell response. Groups of Balb/C mice (n=5 animals/group) were immunized i.m. in each leg with 1 µg repRNA loaded in LNPs and were evaluated for vaccine-elicited immune responses. Shown are counts of total GC B cells (a), counts of follicular helper T cells (b), gating for antigen-specific B cells (c), counts for antigen-specific B cells (d), and the frequency of IgG+ antigen-specific B cells (e). Also displayed are histogram of antigen-specific B cells with equivalent IgG-expression levels (f), and the mean fluorescence intensity (MFI) of IgG+ antigen-specific B cells (g). Serum antibody responses were quantified by ELISA assay conducted on mouse sera collected from 0 to 112 days post-vaccination is shown in (h). Two-way ANOVA with Tukey’s HSD Test was conducted (*, p<0.05; **, p<0.01; ***, p<0.001; ****, p<0.0001); (i) shows endpoint titer at day 14 post-vaccination. Percentage of antigen-specific IgG out of total IgG secreted by bone-marrow plasma cells are shown in (j). Antigen-specific T cell (splenocyte) activity (k-l) were quantified by IFN-γ ELISpot: (k) shows a representative image of the ELISpot wells for each group, and (l) shows number of spots counted per 10^6^ cells in response to overlapping peptides for the HIV immunogen. Each column graph shows means ± s.e.m. Statistics represent one-way ANOVA and Tukey’s HSD Test (ns= not significant; *, p<0.05; **, p<0.01; ***, p<0.001; ****, p<0.0001).

A single immunization with N332-GT2 trimer-encoding repRNA co-delivered with *Ifnar1*-targeting siRNA led to rapid seroconversion, with all animals expressing high titers of trimer-specific IgG by day 14, while mean titers of repRNA alone were ∼2 logs lower at this time point (**Fig. 5h-i**). Antibody titers elicited by repRNA alone slowly rose but even following their plateau at 60 days, they remained 1.4 logs lower than titers in the repRNA/siRNA co-delivery group. repRNA/siIFNAR1 vaccination induced higher levels of IgG2b and a trend toward increased IgG1 compared to repRNA, but other isotypes were elicited at similar levels (**Extended Data Fig. 5**). ELISPOT analysis of trimer-specific and total IgG-producing cells from bone marrow at 6 weeks post-vaccination (**Extended Data Fig. 6a-b**) revealed a trend toward increased antigen-specific plasma cell responses elicited by co-delivery of siIFNAR1 with repRNA in the same LNPs (**Fig. 5j**). Finally, antigen-specific T cell responses in the spleen were also amplified by repRNA/siRNA co-delivery compared to the baseline repAg vaccine or with a scramble siRNA or the co-treatment group, as read out by IFN-γ ELISPOT (**Fig. 5k-l**). Altogether, these results indicate that *Ifnar1* silencing during replicon vaccination can augment diverse elements of the innate and adaptive immune response.

### *Ifnar1* silencing enhances modRNA vaccine-elicited immune responses

We found that empty LNPs used in this study (‘LNP’), as well as clinical formulations used in COVID-19 vaccines from Moderna (mRNA-1273; ‘MDR’) and Pfizer-BioNTech (BNT162b2; ‘PFZ’) induce significant amounts of cytokine production local to the administration site, including type I IFNs (**Extended Data Fig. 7**). To assess whether *Ifnar1* silencing could also benefit non-replicating mRNA vaccines, we first co-transfected siRNA against *Ifnar1* with modRNA encoding for GFP in C2C12 murine myoblasts *in vitro* (**Extended Data Fig. 8**). At 24h post-LNP transfection, we observed that co-delivery of siRNA against *Ifnar1* with modRNA (modGFP/siIFNAR1) increased the expression of GFP over the delivery of modGFP alone (modGFP).

Next, we tested the impact of co-delivering modRNA vaccines with *Ifnar1*-targeted siRNA on vaccine-elicited immune responses *in vivo*. Mice were immunized with LNPs loaded with modRNA encoding N332-GT2 trimer alone (‘(modRNA)’), modRNA co-delivered with siIFNAR1.2 (‘(modRNA/siIFNAR1)’), modRNA co-delivered with scrambled-sequence control siRNA (‘(modRNA/siScramble)’), or with a co-treatment vaccine putting modRNA and siRNA in separate LNPs (‘(modRNA) + (siIFNAR1)’). At 2-weeks post-vaccination, we observed approximately 2-fold increases in the number of GC B cells (**Fig. 6a**) and T follicular helper cells (**Fig. 6b**) in mice vaccinated with the co-delivery vaccine compared to modRNA alone. Antigen-specific GC B cell responses were also significantly increased with siRNA co-delivery, with the total number of antigen-specific GC B cells increasing by 3.6-fold, and the proportion of IgG class-switched cells by ∼9-fold compared to replicon-only group (**Fig. 6c-e**). In GC B cells with similar IgG expression (**Extended Data Fig. 9a**), these antigen-specific B cells also displayed 1.7-fold higher mean fluorescence intensity in binding to antigen tetramers (**Fig. 6f-g**), again suggesting that the B cells generated by the modRNA/siIFNAR1 co-delivery vaccine have higher binding affinity. Lastly, co-delivery of modRNA encoding for the N332-GT2 trimer with siRNA against *Ifnar1* induced rapid seroconversion, such that by week 2 post-vaccination, mice vaccinated with the co-delivery vaccine showed a trend toward ∼1.3 log-fold higher antibody titer (**Fig. 6h** and **Extended Data Fig. 9b**). Thus, *Ifnar1* silencing can also substantially augment non-replicating mRNA vaccines.

**Figure 6.**
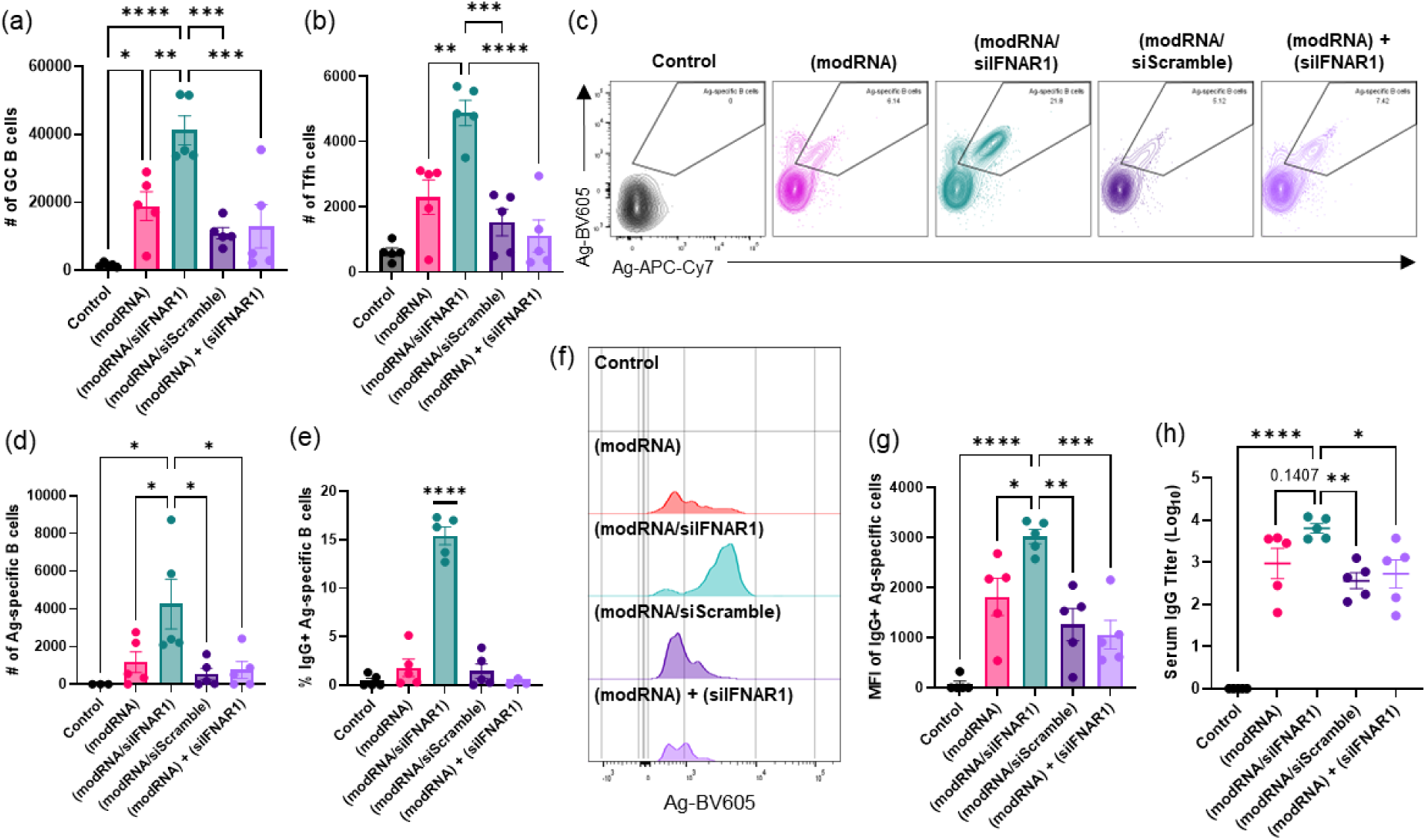
Antigen-specific immunity from modRNA vaccines improve with siRNA co-delivery. Groups of balb/C mice (n=5 animals/group) were immunized i.m. in each leg with 5 µg modRNA loaded in LNPs and were evaluated for vaccine-elicited immune responses. Shown are counts of total GC B cells (a), follicular helper T cells (b), gating for antigen-specific B cells (c), counts for antigen-specific B cells (d), and the frequency of IgG+ antigen-specific B cells (e). Histogram of antigen-specific B cells with equivalent IgG-expression levels is shown in (f), and the mean fluorescence intensity (MFI) of IgG+ antigen-specific B cells are shown in (g). Serum antibody responses were quantified by ELISA assay conducted on mouse sera collected at 14 days post-vaccination and the endpoint titer at 2-weeks post-vaccination are shown in log scale in (h). Each column graph shows means ± s.e.m. Statistics represent one-way ANOVA and Tukey’s HSD Test (*, p<0.05; **, p<0.01; ***, p<0.001; ****, p<0.0001).

### Impact of the LNP composition on siIFNAR1-mediated enhancement of vaccine-elicited immune response

To discern the impact of LNP formulation on the observed improvements in vaccine efficacy with siRNA co-delivery, we finally carried out a vaccination study using the clinical LNP formulation employed in Moderna’s COVID-19 vaccine (mRNA-1273). We immunized Balb/C mice with either repRNA or modRNA encoding N332-GT2 HIV Env trimer, with or without *Ifnar1*-targeting siRNA (**Extended Data Fig. 10**). As we saw with our replicon-tailored LNPs, the Moderna LNP formulation co-loaded with the immunogen-encoding RNA and siIFNAR1 also yielded significant increases in the vaccine-elicited immune response over the modRNA or repRNA vaccine groups without siIFNAR1, including increased total GC B cells (**Extended Data Fig. 10a**), increased Tfh cells (**Extended Data Fig. 10b**), and increased antigen-specific B cells – by 8.7-fold for the modRNA/siIFNAR1 vaccine and 11.3-fold for repRNA/siIFNAR1 (**Extended Data Fig. 10c-d**). The proportion of IgG class-switched antigen-binding GC B cells also increased by 10-fold for modRNA/siIFNAR1 vaccine and ∼5-fold for repRNA/siIFNAR1 (**Extended Data Fig. 10e**). As seen with our replicon-tailored LNPs, the mean fluorescence intensity of trimer-binding IgG^+^ B cells also increased by ∼2.5-fold for both modRNA/siIFNAR1 and repRNA/siIFNAR1 vaccine groups (**Extended Data Fig. 10f-g**). Finally, we observed significant increases in the serum antibody titer in the siIFNAR1 co-delivery vaccine over the modRNA or repRNA vaccine alone (**Extended Data Fig. 10h-i**). These results suggest that siIFNAR1 co-delivery may be a generalizable strategy to improve vaccine-elicited immune response of repRNA and modRNA vaccines.

## DISCUSSION

Alphavirus-derived replicons are of interest as a platform for RNA-based gene delivery as their self-amplification in transfected cells can enable very high, sustained levels of therapeutic payload production compared to conventional mRNAs.^40,41^ In the setting of vaccines, these features are of interest because the duration of antigen availability can directly impact key features of the immune response such as germinal center development^42,43^ and T cell priming^44^, and repRNA may also provide practical advantages such as enabling lower total doses of RNA (and its inflammatory LNP carrier) to be used. Progress is being made in the clinical testing of repRNA-based vaccines^45–47^, but early studies have shown that achieving the potential benefits of replicons may require multiple levels of optimization^6,48^. A key factor is the interaction between repRNA and innate immune sensing pathways, which can both promote inflammatory “danger signals” that promote the immune response elicited by repRNA vaccines, but also activate translational inhibition mediated by the antiviral response that blunts the development of adaptive immunity.

Prior studies examining the roles of innate immune sensing in repRNA function identified that type I IFN plays a major role in inhibiting replicon expression, but entirely eliminating this pathway using mouse models genetically deficient in type I IFN sensing have reported mixed results on vaccine immunogenicity. The evaluation of repRNA responses in IFN-knockout animals is complicated by the many roles type I interferon plays in the natural immune response, such as promoting immune cell recruitment to the injection site, stimulating DCs, supporting differentiation of CD4^+^ and CD8^+^ effector T cells and Tfh cells, ultimately leading to robust B cell differentiation into antibody-secreting plasma cells.^26,27,29,49,50^ We hypothesized that formulation of LNPs carrying repRNA together with siRNA against *Ifnar1* could focus type I IFN signaling inhibition on transfected cells and limit the duration of IFN modulation, to avoid blocking potentially beneficial IFN signaling in responding immune and stromal cells in the injection site and/or draining lymph nodes. In mice, we found that 3 days of robust *Ifnar1* silencing led to drastically enhanced replication and expression of repRNA by over 10-fold at both transcriptomic and proteomic levels, correlating with increased antigen-specific B (∼8-fold higher antigen-specific GC B cells) and T (4.4-fold) cell responses, as well as >10-fold increased serum antibody titers compared to vaccines delivered without any siRNA or with scramble sequence of siRNA against *Ifnar1*. We found similar enhancements in the immunogenicity of non-replicating RNA vaccines: While modRNA is designed to evade innate recognition, emerging evidence suggests that modRNA is not completely immune-silent *in vivo*, and the LNP delivery vehicle is known to stimulate type I IFN production.^18–20^ Thus, this approach appears generally useful for RNA-based vaccines. Notably, we found that it is key to encapsulate antigen-encoding RNAs and siRNA against *Ifnar1* in the same LNP for co-delivery into the same cells, as opposed to independent delivery of the two RNA payloads in separate LNPs to the same tissue, but not necessarily the same cell.

Several prior efforts have been made to suppress type 1 IFN activation with regard to improving RNA transcript expression. In one instance, innate immunity was suppressed by co-delivering the repRNA with mRNA encoding for vaccinia virus immune evasion proteins, such as E3 and B18. E3 displayed superior blockade of protein kinase R (a cytoplasmic RNA sensor) and B18 effectively inhibited OAS1 (anti-viral enzyme that degrades viral RNA) for an overall downregulation of interferon-β and enhanced repRNA expression *in vivo*.^51^ In similar fashion, innate inhibiting proteins (IIPs) have also been co-encoded into the replicon to augment the primary cargo expression.^12^ IIPs, specifically PIV-5 V and MERS-CoV ORF4a proteins, enhanced protein expression by 100-to 500-fold *in vitro* in IFN-competent cells and abated dose nonlinearity *in vivo*, which led to a 10-fold increase in the rabies virus-specific IgG titer and neutralization IC_50_ values in rabbits. However, the effects were lost when tested in mice or rats. Both of these strategies also introduce a second immunogenic payload to be expressed, which raises concerns for potential appropriation of immunodominance from the primary antigen.

Another strategy to further enhance the activity and immunogenicity of repRNA vaccines is to substitute modified nucleotides into the sequence.^52^ As demonstrated with conventional mRNA used in the clinically approved COVID-19 vaccines, modified nucleotides, such as N1-methylpseudouridine (N1mΨ), are able to improve RNA stability and limit innate immune recognition. Most modified nucleotide substitutions are generally thought to be incompatible with repRNA and render them dysfunctional^53–56^, or to have no impact on repRNA activity, as the repRNA is amplified in cells with native unmodified nucleotides^57,58^. However, recent work identified certain modified nucleotides that are tolerated by repRNAs and demonstrated that substitution with 5-methylcytidine nucleotides in a SARS-CoV-2 vaccine is able to decrease IFN-α production by ∼2.5-fold and improve survival against viral challenge *in vivo–* at least under the condition of very low doses of replicon RNA (10 ng). This approach may be synergistic with the strategy taken here.

The most comprehensive work done to understand the role of type 1 IFN in RNA vaccination suggests that the timing and intensity of type 1 IFN induction relative to the activation of T cell receptors (TCRs) determines whether the innate response inhibits or stimulates vaccines.^59^ These studies found that local administration of RNA *via* subcutaneous^24,25^, intradermal^24,30^, or intramuscular^11,12^ routes triggers production of type 1 IFNs at the site of injection, activates dendritic cells (DCs) to carry and present the processed antigen to T cells in the draining lymph nodes to activate TCRs and IFNAR1, leading to a delayed pro-apoptotic and anti-proliferative transcription program. On the other hand, systemic administration^34,60^ of RNA allows localization in the spleen, where DCs can directly express the antigen for immediate TCR and IFNAR1 activation on T cells to promote T cell proliferation, differentiation, and survival.

Overall, our findings demonstrate that while innate recognition of repRNA and LNPs can inhibit RNA vaccine function, transiently suppressing the type 1 IFN response both augments antigen expression and enhances early injection site and lymph node innate immune activation, leading to dramatically improved downstream vaccine-elicited immune responses. *Ifnar1* silencing through siRNA co-delivery appears to be a simple approach to augment many elements of vaccine-elicited immunity for both replicating and non-replicating RNA vaccines, which could be readily translated to humans.

## ONLINE METHODS

### Materials

*N*^1^,*N*^3^,*N*^5^-tris(3-(didodecylamino)propyl)benzene-1,3,5-tricarboxamide (TT3) was synthesized as previously described^37^; (6*Z*,9*Z*,28*Z*,31*Z*)-Heptatriaconta-6,9,28,31-tetraen-19-yl 4-(dimethylamino) butanoate (DLin-MC3-DMA) was purchased from MedChemExpress (CAT#HY-112251); 1,2-dioleoyl-*sn*-glycero-3-phosphoethanolamine (DOPE; CAT#850725), Cholesterol (CAT#700100), 1,2-dimyristoyl-*rac*-glycero-3-methoxypolyethylene glycol-2000 (DMG-PEG2k; CAT#88015) were purchased from Avanti Polar Lipids. 1,1’-Dioctadecyl-3,3,3’,3’-Tetramethylindodicarbocyanine, 4-Chlorobenzenesulfonate Salt (DiD; CAT#D7757) was purchased from ThermoFisher Scientific. Soluble HIV Env trimer protein N332-GT2 was prepared as previously described.^35^ Citrate buffer (pH 3; CAT#J61391-AK) was purchased from Alfa Aesar. For dialysis, 20K MWCO Slide-A-Lyzer™ MINI Dialysis Device (ThermoFisher Scientific), RNAse-free PBS (AM9625; ThermoFisher Scientific) was used. Quant-iT RiboGreen RNA kit (CAT#R11490, Invitrogen) was used to evaluate the RNA encapsulation efficiency in LNPs. Firefly luciferase-encoding modRNA (CleanCap® FLuc mRNA (5moU); CAT# L-7202) was purchased from Trilink Biotechnologies. For bioluminescence studies, luciferin was purchased from GoldBio (CAT# LUCK-1G).

### RNA synthesis

Development of a backbone for self-amplifying RNAs (repRNA) was described previously.^61–63^ In brief, structural genes from the Venezuelan Equine Encephalitis Virus genome were replaced with a reporter gene and favorable mutations were introduced via *in vitro* evolution. In this study, we replaced the reporter gene of this backbone with green fluorescent protein (GFP), mCherry, firefly luciferase, or HIV immunogen N332-GT2 (a transmembrane form of a stabilized SOSIP HIV envelope trimer^35^) using traditional cloning methods. For modRNA Synthesis, template DNA plasmids used in the production of modRNA were created using a commercially available Cloning Kit for mRNA Templates (Takara #6143) according to manufacturer’s instructions. Resultant plasmid DNA was linearized *via* endonuclease digestion and purified with PureLink PCR Purification columns (ThermoFisher #K310002) following the manufacturer’s instructions. To synthesize RNA, 20 μL *in vitro* transcription (IVT) reactions were performed using the HiScribe T7 High Yield RNA Synthesis Kit (NEB #E2040S) and 1-2 μg of linear DNA template (scaled as needed). For modRNA, modified base N1-methylpseudouridine triphosphate (TriLink #N-1081) was added to the reaction mixture instead of canonical uridine triphosphate, and CleanCap Reagent AG (TriLink #N-7113) was utilized to co-enzymatically add 5’ Cap-1 structures to synthesized RNA. The IVT product was purified using PureLink RNA Mini columns (ThermoFisher #12183018A) following the manufacturer’s instructions. Replicon RNA was capped and methylated using the ScriptCap Cap 1 Capping System (CellScript #C-SCCS2250) following the manufacturer’s instructions, after which RNA was purified a final time using PureLink RNA Mini columns. Quality of the resulting repRNA was assessed using UV-Vis spectrophotometry and gel electrophoresis.

### Lipid nanoparticle synthesis

Lipids were stored in ethanol at -20°C, and RNA constructs were stored in RNAse-free water at - 80°C and were thawed on ice before use. Lipid nanoparticles (LNPs) were synthesized using a microfluidic organic-aqueous nanoprecipitation method, as previously described.^64^ The organic phase was prepared by solubilizing the lipids TT3^37^, Dlin-MC3-DMA, DOPE, Cholesterol, and DMG-PEG2k in ethanol at a molar ratio of 10:25:20:40:5. For fluorescently labeled LNPs, DiD was added to the organic phase at 0.1 mol%. The aqueous phase of RNA was prepared by diluting the RNA (stored in RNAse-free water) with 10 mM citrate buffer, pH 3.0 (CAT#J61391-AK; Alfa Aesar). For empty LNP synthesis, RNA was replaced with additional citrate buffer. The two phases were prepared at an ethanol:aqueous volume ratio of 1:2 (for mRNA, volume ratio was 1:3), and RNA and lipids combined at an N:P (number of amines on TT3 and MC3 to number of phosphates on repRNA) ratio of 3.7:1 (for mRNA, N:P ratio was set to 8:1). Each phase was loaded into a syringe (BD), and locked onto the NxGen microfluidic cartridge for mixing using a NanoAssemblr Ignite instrument (Precision Nanosystems). The Ignite was set to operate with the following settings: volume ratio-2:1 (for mRNA, 3:1); flow rate-12 ml/min; waste volume-0 mL. The resulting LNPs were dialyzed against PBS using 20K MWCO Slide-A-Lyzer™ MINI Dialysis casettes (ThermoFisher Scientific) at 25°C for 90 min, with an exchange of the buffer reservoir after 45 min. Other LNP formulations were synthesized in the same manner with different lipid compositions according to **Supplementary Table S1**.

### Particle characterization

Hydrodynamic size, polydispersity index, and zeta-potential of the LNPs was determined by dynamic light scattering (DLS; Malvern Panalytical). To measure hydrodynamic size, 10 uL of particles were diluted in 800 uL of deionized water and placed into a 1.5 mL cuvette (Fisher Scientific) and measured. Same sample preparation was used to measure zeta-potential using the folded capillary zeta cell (Malvern Panalytical). Cryogenic electron microscopy (cryo-EM) was used to qualitatively assess the LNP size and polydispersity as previously described.^64^ Briefly, 3 uL of the particles in deionized water was dropped on a lacey copper grid coated with a continuous carbon film. Excess liquid was blotted and the grid was frozen in liquid ethane cooled by liquid nitrogen in the Gatan Cryo Plunge III. Then the grid was mounted on a Gatan 626 single tilt cryo-holder of the TEM column while being kept chill with liquid nitrogen. Imaging on a JEOL 2100 FEG microscope was operated at 200 kV with a magnification range of 10,000–60,000. Gatan 2kx2k UltraScan CCD camera was used to record all images.

Quant-iT RiboGreen RNA kit (Invitrogen) was used to evaluate the RNA encapsulation efficiency in LNPs according to the manufacturer instructions. In a black 96 well plate, control RNA provided in the kit was used to make two sets of standard curves with serial dilution in either RNAse-free water or Triton-X-100. Samples were 100-fold diluted in either RNAse-free water (to quantify RNA present outside of LNPs) or Triton-X-100 (to quantify total RNA present inside and outside of LNPs). Fluorescence was measured with excitation wavelength= 480 nm, and emission wavelength= 520 nm.

### *In vitro* cell culture and transfection

C2C12 mouse myoblasts were cultured in growth media (DMEM supplemented with 10% FBS and 1% penicillin/streptomycin, and were incubated at 37°C in 5% CO_2_). For transfection, 25,000 cells were seeded in 96 well plates for overnight attachment, and 1 ug of GFP-encoding repRNA loaded in DiD-labeled LNPs in 10 uL of PBS were added with 20 uL Opti-MEM (Gibco) and 70 uL of growth media. At 24h post-transfection, cells stained with Zombie Aqua (BioLegend) for cell viability, and were transferred to a V-bottom 96 well plate for quantification of GFP expression on the BD LSRFortessa™ Cell Analyzer (BD Biosciences).

### Mice

All animal studies and procedures were carried out following federal, state, and local guidelines under an IACUC-approved animal protocol. Female Balb/c (JAX Stock No. 000651) mice, C57Bl/6 (JAX Stock No. 000664), and B6(Cg)-Ifnar1tm1.2Ees/J (JAX Stock No. 028288) at 6–8 weeks of age were maintained in the animal facility at the Massachusetts Institute of Technology (MIT). For all LNP administrations, mice were intramuscularly injected in the left and right gastrocnemius muscles at dose of 1 µg repRNA or 5 µg mRNA in 20 µL of PBS.

### *In vivo* mouse imaging

To assess the efficacy of different LNP formulations using repRNA or mRNA encoding for firefly luciferase as a reporter, we immunized Balb/c mice intramuscularly (i.m.) with 1 ug repRNA or 5 ug mRNA doses of LNPs in each of the left and right gastrocnemius muscles. At days 1, 3, 7, 10, and 15 post-i.m. injection, the mice were intraperitoneally (i.p.) administered 200 µL of luciferin (50 mg/ml in PBS), and imaged using the In Vivo Imaging System (Xenogen IVIS 200; PerkinElmer) 10 min post-i.p. injection.

### qRT-PCR for relative quantification of RNA expression

For *in vitro* quantification of RNA expression, C2C12 mouse myoblasts were seeded at 300,000 cells/well in a 6 well-plate and incubated overnight for attachment. repRNA encoding for GFP was loaded in LNPs and treated at 5 ug/well in 100 uL of PBS, 200 uL of Opti-MEM and 700 uL of growth media for overnight transfection. Total RNA was purified from the cells using a PureLink™ RNA Mini Kit (Invitrogen). Briefly, cells were lysed and the lysates underwent a series of fast spin-column washes. The final RNA concentration was measured using a NanoDrop UV-VIS spectrometer (ThermoFisher Scientific). The relative quantification of RNA expression was assessed using a two-step quantitative real-time reverse transcription polymerase chain reaction (qRT-PCR, QuantStudio, ThermoFisher). cDNA was synthesized from RNA using the iScript™ Reverse Transcription Supermix (Bio-Rad), and PCR was run with the iTaq™ Universal SYBR® Green Supermix (Bio-Rad). Primers used are listed in **Supplementary Table S3**.

For *in vivo* quantification of RNA expression, mice were intramuscularly administered 1 ug of repRNA encoding for GFP loaded in LNPs. At days 1, 3, 7, and 14 post-administration, mice were sacrificed and the gastrocnemius and popliteal lymph nodes were harvested. Tissues were kept in RNAlater™ Stabilization Solution (Invitrogen) at -20°C until thawed for RNA purification. Tissues were placed in 1 mL of Trizol (Invitrogen) with β-mercaptoethanol (Gibco), and were homogenized in gentleMACS™M tubes using gentleMACS™ Octo Dissociator (Miltenyi Biotec) and its built-in muscle protocol at least 5 times or until no large tissue fragments was visible. The M tubes were centrifuged at 600 x g for 5 min at 4°C. Supernatants were collected and gently mixed with chloroform, which was added at 20% by volume and incubated on ice for 2-3 min. Samples were then centrifuged for 15 minutes at 12,000 × g at 4°C to phase-separate the mixture. The top aqueous phase was transferred to new RNAse-free tubes, and 500 uL of isopropanol was added for 10 min incubation at 4°C. RNA was pelleted by centrifugation for 10 minutes at 12,000 × g at 4°C. The pellet was resuspended and washed in 1 mL of 70% ethanol by centrifugation for 5 minutes at 7500 × g at 4°C. The pellet was dried briefly in air for 5 min before being resuspended in 50 uL of RNAse-free water, and placed on a heat block for 10 min at 60°C. RNA concentration was measured by NanoDrop and were stored in -80°C until cDNA synthesis using iScript™ Reverse Transcription Supermix and qRT-PCR with iTaq™ Universal SYBR® Green Supermix. Primers used are listed in the **Supplementary Table S3**.

### Cryofluorescence Tomography

Mice were placed on alfalfa-free diet to minimize autofluorescence from the gut for a week prior to repRNA administration. Mice were intramuscularly administered 1 ug of repRNA encoding for mCherry loaded in LNPs in each leg, with 10% of the LNPs labeled with DiD (ThermoFisher). At days 1, 3, and 7 post-repRNA administration, mice were euthanized by CO_2_, and skin was removed to eliminate autofluorescence. A holder made of polylactic acid (PLA) was 3D-printed to control orientation of the mouse body during freezing in a hexane bath at dry ice temperature for 5 minutes. Afterwards, the frozen samples were placed in a -20°C freezer overnight to degas the hexane. Mouse lower limbs were isolated from the holder and were mounted in an OCT “C” block (14×10.5×7cm) and sliced and imaged using an EMIT (Baltimore, MD) Xerra CFT system. Slices were cut at 35 µm thicknesses and images were acquired using excitation wavelength at 555 nm and detection filter of 586/15 nm for mCherry and excitation wavelength at 640 nm and detection filter of 680/13 nm for DiD detection. Images were analyzed using Fiji with Bio-formats.

### mCherry quantification from muscles

Mice were intramuscularly administered 1 µg of repRNA encoding for mCherry loaded in LNPs. At day 7 post-administration, mice were sacrificed and the left and right gastrocnemius muscles were harvested. Muscles were placed in gentleMACS™M tubes (Miltenyi Biotec) with 300 uL of lysis buffer composed of RIPA buffer (ThermoFisher Scientific) supplemented with 1% EDTA (Invitrogen) and 1% Halt™ Protease and Phosphatase Inhibitor Cocktail (ThermoFisher Scientific). Muscles were lysed using gentleMACS™ Octo Dissociator (Miltenyi Biotec) and its built-in muscle protocol at least 5 times or until no large tissue fragments was visible. The M tubes were centrifuged at 600 x g for 5 min at 4°C. Supernatants were collected and stored in -20°C until use. For protein quantification, mCherry ELISA kit (CAT# ab221829; Abcam) was used according to manufacturer instructions. Briefly, a standard curve was set using the provided mCherry recombinant protein with serial dilution from 4,000 pg/mL with dilution factor of 2x. Muscle lysates were thawed and 200-fold diluted in sample diluent buffer in each well. In each well, 50 uL of standard or sample and 50 uL of the provided antibody cocktail were placed. The plate was covered and incubated for 1h at 25°C on a shaker. After three consecutive washes with the provided wash buffer, 100 uL of TMB development solution was added to each well and incubated for 1 min. Finally, 100 uL of the provided stop solution was added and the absorbance was measured at 450 nm using a UV-Vis plate reader.

### Mouse muscle and lymph node immunophenotyping

Mice were intramuscularly administered 1 µg of repRNA encoding for either the antigen or mCherry loaded in LNPs. At days 3 and/or 7 post-administration, mice were sacrificed and the left and right gastrocnemius muscles and popliteal lymph nodes were harvested. Tissues were placed in gentleMACS™C tubes (Miltenyi Biotec) with 5 uL of digestion buffer composed of complete RPMI + 0.1% w/v Collagenase I + 0.1% w/v Collagenase IV. Muscles were lysed using gentleMACS™ Octo Dissociator (Miltenyi Biotec) and its built-in muscle protocol at least 2 times or until no large tissue fragments was visible. The samples were incubated at 37°C for 1h on shaker for further digestion. Post-digestion, samples were filtered through 70 µm pores twice and centrifuged at 600 x g for 5 min at 4°C to isolate pellets composed of single cells. Zombie Aqua (BioLegend) in PBS was used to stain for cell viability, and antibody staining was performed in fluorescence-activated cell sorting (FACS) buffer (PBS, 1% bovine serum albumin (BSA), 0.02% NaN3, and 2 mM EDTA). Fc-mediated binding was blocked using purified anti-CD16/32 (2.5 μg/ml; 93; BioLegend) at 4°C for 15 min, on top of which we added primary antibodies for cell surface staining at 4°C for 30 min. Immune cells were identified with anti-CD45 PerCP-Cy5.5 (30-F11; BioLegend), B cells were stained with anti-B220 BV421 (RA3-6B2; BioLegend) and anti-CD19 PE-Cy7 (6D5; BioLegend), T cells were marked by anti-CD3 BUV661 (145-2C11; BD Biosciences), macrophages were stained with anti-F4/80 APC-Cy7 (BM8; BioLegend) and anti-CD11b BUV396 (M1/70; BioLegend), monocytes were stained with anti-Ly6C BV650 (HK1.4; BioLegend), neutrophils with anti-Ly6G APC (1A8; BioLegend), and anti-CD11b. Dendritic cells were stained with anti-CD11c AF488 (N418; BioLegend), anti-MHC-II AF700 (M5/114.15.2; BioLegend), and anti-CD103 BV786 (M290; BD Biosciences). Activation states of myeloid cells were assessed by staining for anti-CD80 BUV737 (16-10A1; BD Biosciences) and anti-CD86 BV605 (GL-1; BioLegend). Stained cells were fixed with 4% paraformaldehyde (ThermoFisher Scientific) for 10 min at 25°C, washed and resuspended in FACS buffer for flow cytometric analysis on a BD FACSymphony™ A3 Cell Analyzer (BD Biosciences). CountBright™ Absolute Counting Beads (ThermoFisher Scientific) were added to each well immediately before running flow cytometry. Gating strategy is shown in **Supplementary Fig. S1**.

### Germinal center (GC) analysis

Balb/C mice were immunized as described above, and at 2 weeks post-vaccination, popliteal lymph nodes were collected and mechanically dissociated to obtain single cell suspensions. Zombie Aqua (BioLegend) in PBS was used to stain for cell viability, and antibody staining was performed in FACS buffer. Fc-mediated binding was blocked using purified anti-CD16/32 (2.5 μg/ml; 93; BioLegend) at 4°C for 15 min, on top of which we added primary antibodies for cell surface staining at 4°C for 30 min. GC B cells were stained using anti-GL7 PerCPCy5.5 (GL7; BioLegend), anti-CD38 AF488 (90; BioLegend), and anti-B220 PECy7 (RA3-6B2; BioLegend). IgG-expression was characterized by anti-IgG BV786 (X56; BD Biosciences), Follicular helper T cells were assessed using anti-CD4-BV711 (GK1.5; BioLegend), anti-CXCR5-PE (phycoerythrin) (L138D7; BioLegend), and anti-PD1-BV421 (29F.1A12; BioLegend). The HIV env trimer used as the antigen was conjugated to either BV605 (streptavidin-conjugated; BioLegend) or APC-Cy7 (streptavidin-conjugated; BioLegend), and both probes were used to detect antigen-specific B cells. Stained cells were fixed with 4% paraformaldehyde (ThermoFisher Scientific) for 10 min at 25°C, washed and resuspended in FACS buffer for flow cytometric analysis on a BD FACSymphony™ A3 Cell Analyzer (BD Biosciences). CountBright™ Absolute Counting Beads (ThermoFisher Scientific) were added to each well immediately before running flow cytometry. Gating strategy is shown in **Supplementary Fig. S2**.

### Serum antibody titer quantification

We vaccinated healthy Balb/C mice by i.m. injecting 1 µg repRNA or 5 µg mRNA doses of the LNPs in each of the left and right gastrocnemius muscle. At different time points post-i.m. injection, the mice underwent retro-orbital bleeding; blood was collected in Z-gel PP tubes for blood serum collection (CAT#41.1500.005; Sarstedt). Serum was collected by centrifuging blood at 10,000 x *g* for 4 minutes, and stored at -80°C prior to use. To conduct ELISAs, NUNC MaxiSorp plates were coated overnight with 1 µg/mL purified HIV antigen in PBS, then blocked for 2 h with 1% BSA in PBS. Mouse sera were initially diluted 50x in blocking buffer, followed by 3x serial dilutions. Diluted sera were transferred to blocked plates and incubated for 2 h. HRP-conjugated immunoglobulins (e.g., IgG, IgG1, IgG2a, IgG2b, IgG3, IgM; Bio-Rad) were used as detection antibodies at 1:5,000 for endpoint titer assessments, with gp120-specific monoclonal antibody VRC01 used as a positive control. 3,3’,5,5’-Tetramethylbenzidine (TMB) signal was read using a microplate reader by subtracting the absorbance at 450 nm by that at 550 nm.

### Enzyme-linked immunosorbent spot (ELISpot) assay of splenocytes

Spleens were harvested 2 weeks after mice had been vaccinated i.m. with LNPs carrying repRNA encoding for the HIV immunogen in the left and right gastrocnemius muscles of mice. Splenocytes were isolated by mechanical dissociation of the spleen and erythrocytes were removed using the Gibco Ammonium-Chloride-Potassium lysing buffer (Thermo Fisher Scientific). ELISpot was conducted using the mouse IFN-γ ELISPOT Kit (BD Biosciences). Cells were seeded on IFN-γ-coated wells at 10^6^ cells/well in triplicate for pools of overlapping peptides covering the entire sequence of the HIV env trimer (15-mer peptides overlapping by 11 amino acids), which were added to the cells at 2 µg/mL. Cells were stimulated by wrapping the plate in foil and incubating overnight at 37°C. Plates were developed and detected according to manufacturer’s instructions. Plates were scanned using a CTL-ImmunoSpot Plate Reader, and data were analyzed using CTL ImmunoSpot Software.

### Enzyme-linked immunosorbent spot (ELISpot) assay of bone-marrow plasma cells

Balb/C mice were immunized as described above, and at 6 weeks post-vaccination, bone-marrow plasma cells (BMPCs) were isolated from the femur and tibia. ELISpot Flex: Mouse IgG (HRP) kit (CAT# 3825-2H; MABTECH) was used to evaluate antigen-specificity of BMPCs according to manufacturer’s instructions. Briefly, a MSIP PVDF-membrane plate was activated by ethanol washing, then coated with the provided anti-IgG antibody overnight at 4°C. Coated plates were seeded with 10^6^ BMPCs/well for total IgG quantification and 5×10^6^ BMPCs/well for antigen-specific IgG quantification for overnight incubation at 37°C in 5% CO_2_. Cells were removed by washing before target analytes were added to wells; anti-IgG-biotin was added for total IgG quantification, and biotinylated HIV env trimer was added for antigen-specific IgG quantification. Plates were developed and detected according to manufacturer’s instructions. Plates were scanned using the CTL-ImmunoSpot Plate Reader, and data were analyzed using CTL ImmunoSpot Software.

### Bead-based ELISA cytokine quantification

At 24 h post-i.m. administration of LNPs, gastrocnemius muscles and popliteal lymph nodes were collected from mice. Tissues were placed in 0.3 mL of lysis buffer, composed of RIPA buffer (ThermoFisher Scientific) supplemented with 1% EDTA (Invitrogen) and 1% Halt™ Protease and Phosphatase Inhibitor Cocktail (ThermoFisher Scientific). Tissues were lysed in gentleMACS™M tubes using gentleMACS™ Octo Dissociator (Miltenyi Biotec) and its built-in muscle protocol at least 5 times or until no large tissue fragments was visible. The M tubes were centrifuged at 600 x g for 5 min at 4°C. Supernatants were collected and stored in -20°C until use. The lysates from mouse gastrocnemius muscles and popliteal lymph nodes were analyzed using the Legendplex mouse antivirus response panel (Biolegend) following the manufacturer’s suggested protocol and analyzed using the LEGENDplex Data Analysis Software Suite.

## ACKNOWLEDGEMENTS

We thank the Koch Institute’s Robert A. Swanson (1969) Biotechnology Center for technical support, specifically the Flow Cytometry Core, the Peterson (1957) Nanotechnology Materials Core Facility, and the Preclinical Modeling, Imaging & Testing Core (PMIT). Figure schematics were created in BioRender.com. This work was supported by the NIH (awards AI176533 and UM1AI144462 to DJI, R35GM144117 to YD, and CA265706 to DJI and YD), the Marble Center for Nanomedicine, and the Ragon Institute of MGH, MIT, and Harvard. This work was also supported in part by the Koch Institute Support (core) Grant P30-CA014051 from the National Cancer Institute. DJI is an investigator of the Howard Hughes Medical Institute.

## AUTHOR CONTRIBUTIONS

Conceptualization, BJK and DJI; Methodology, BJK, RRH, HM; Investigation, BJK, RRH, TR, JD, HM, JYJ, MCB, WA, LM, MBM, BL, YZ; Writing-original draft, BJK and DJI; Writing-review and editing, all authors; Resources and Supervision, YD and DJI.

## EXTENDED DATA

**Extended Data Figure 1.**
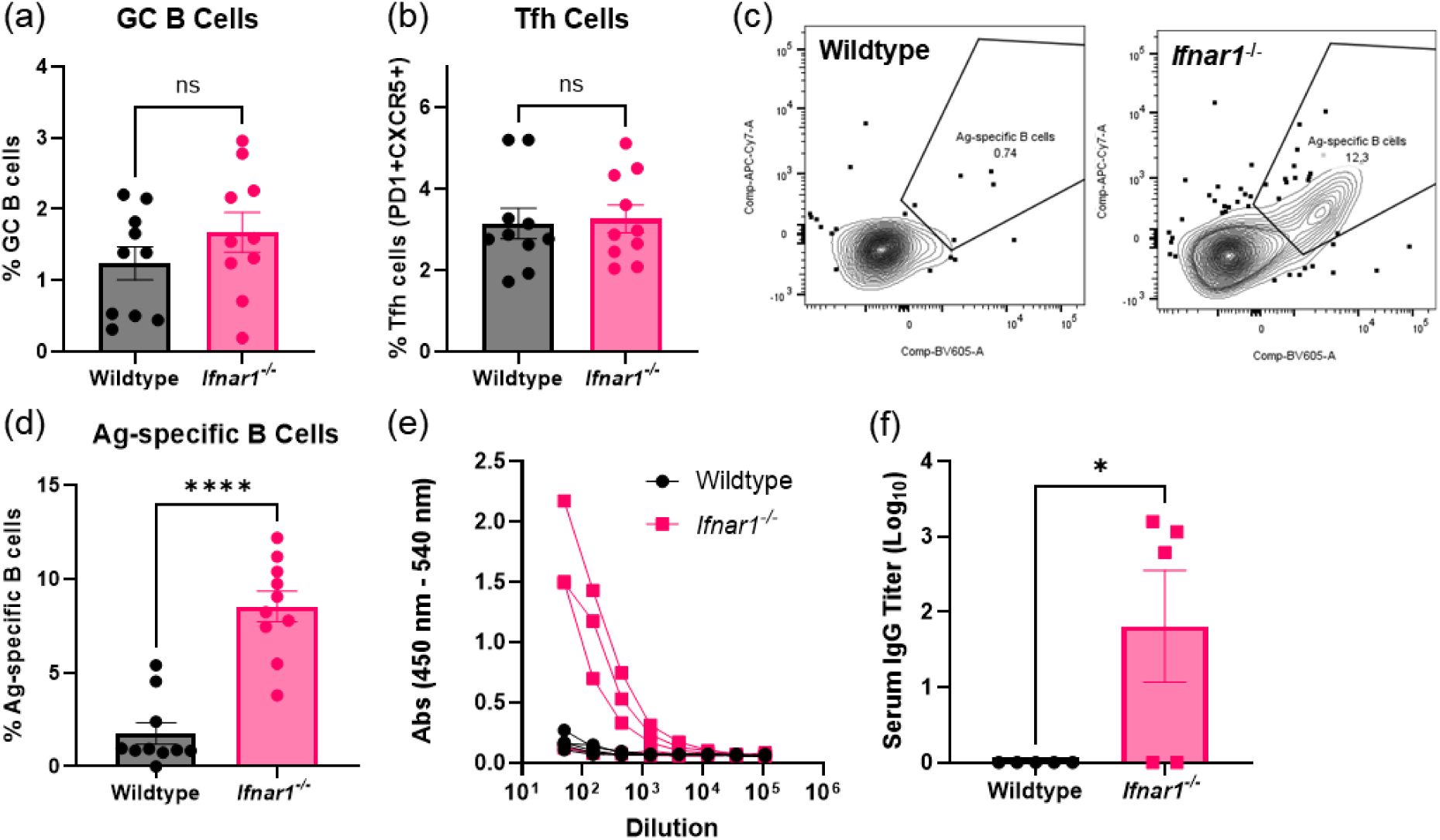
Impact of IFNAR1 on germinal center (GC) responses and antibody-production. Groups of C57Bl/6 (Wildtype) and B6(Cg)-Ifnar1tm1.2Ees/J (*Ifnar1*^-/-^) mice (n=5 animals/strain) were injected i.m. in both the left and right gastrocnemius muscles with 1 µg replicon RNA (encoding for HIV env trimer immunogen) in LNPs. Shown are frequencies of germinal center B cells (a) and follicular helper T (Tfh) cells (b), flow plots of antigen-specific B cells in Wildtype and *Ifnar1*^-/-^ mice (c), frequencies of antigen-specific B cells (d), individual dilution curves of serum antibody titer for each mouse (e), and the endpoint titer (f). means ± standard error means and statistics represent one-way ANOVA and Tukey’s HSD Test (ns= not significant, *, p<0.05; ****, p<0.0001).

**Extended Data Figure 2.**
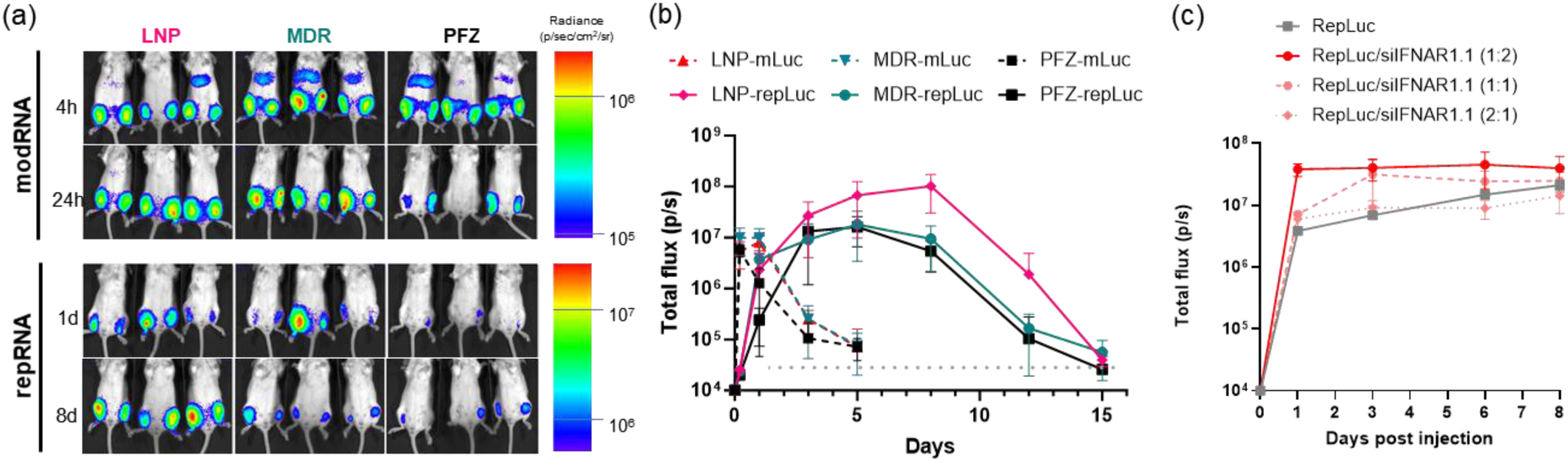
Lipid nanoparticle screening and validation for clinical potential. Groups Balb/c mice (n = 3 animals/group averaged across a total of 6 legs/group) were injected i.m. in both the left and right gastrocnemius muscles with 1 µg repRNA or 5 µg modRNA (encoding firefly luciferase) in LNPs in comparison against clinically used formulations (MDR: Moderna, PFZ: Pfizer). Shown are luciferase reporter signals over time; representative photograph/false-color overlays are shown in (a), and luciferase reporter signals over time are shown in (b); dotted grey line indicates background signal of untreated mice, shown are means ± standard deviation (n=6). (c) Groups Balb/c mice (n = 3 animals/group averaged across a total of 6 legs/group) were injected i.m. in both the left and right gastrocnemius muscles with 1 µg replicon RNA encoding for firefly luciferase (RepLuc) with varying mass ratios of siRNA against *Ifnar1* (siIFNAR1) in LNPs. Shown are luciferase reporter signals over time and bars represent means ± standard error means (n=6).

**Extended Data Figure 3.**
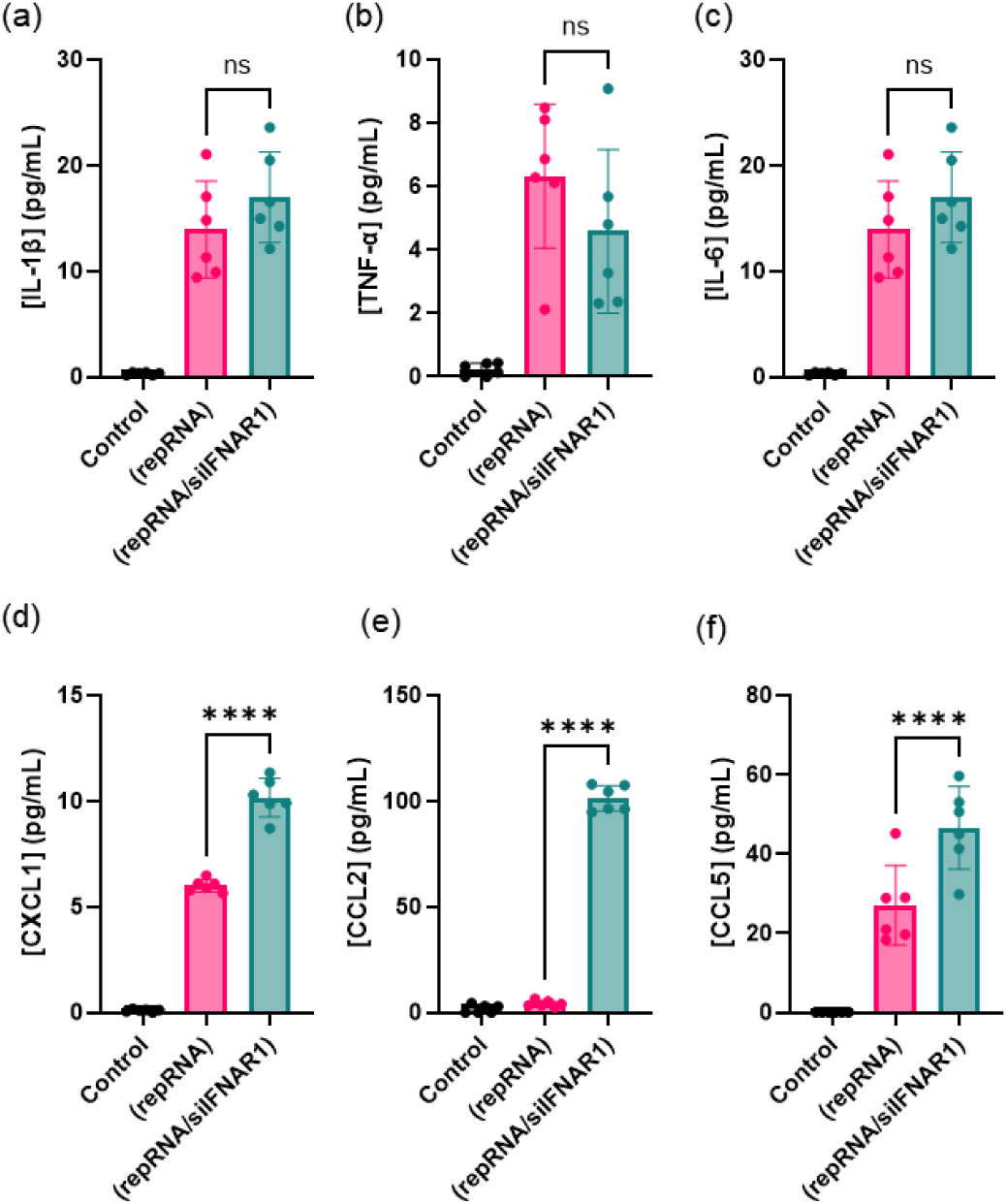
Cytokine profile in the muscle from replicon vaccines. Groups of Balb/C mice (n=6) were immunized i.m. in each leg with LNPs loaded with repRNA encoding for antigen with or without siIFNAR1. At day 3 post-vaccination, gastrocs were harvested and evaluated for cytokine induction. Shown from the muscle are: (a) IL-1β; (b) TNF-α; (c) IL-6; (d) CXCL1; (e) CCL2; and (f) CCL5. Graphs show means ± s.e.m. Statistics represent one-way ANOVA and Tukey’s HSD Test (ns, not significant; ****, p<0.0001).

**Extended Data Figure 4.**
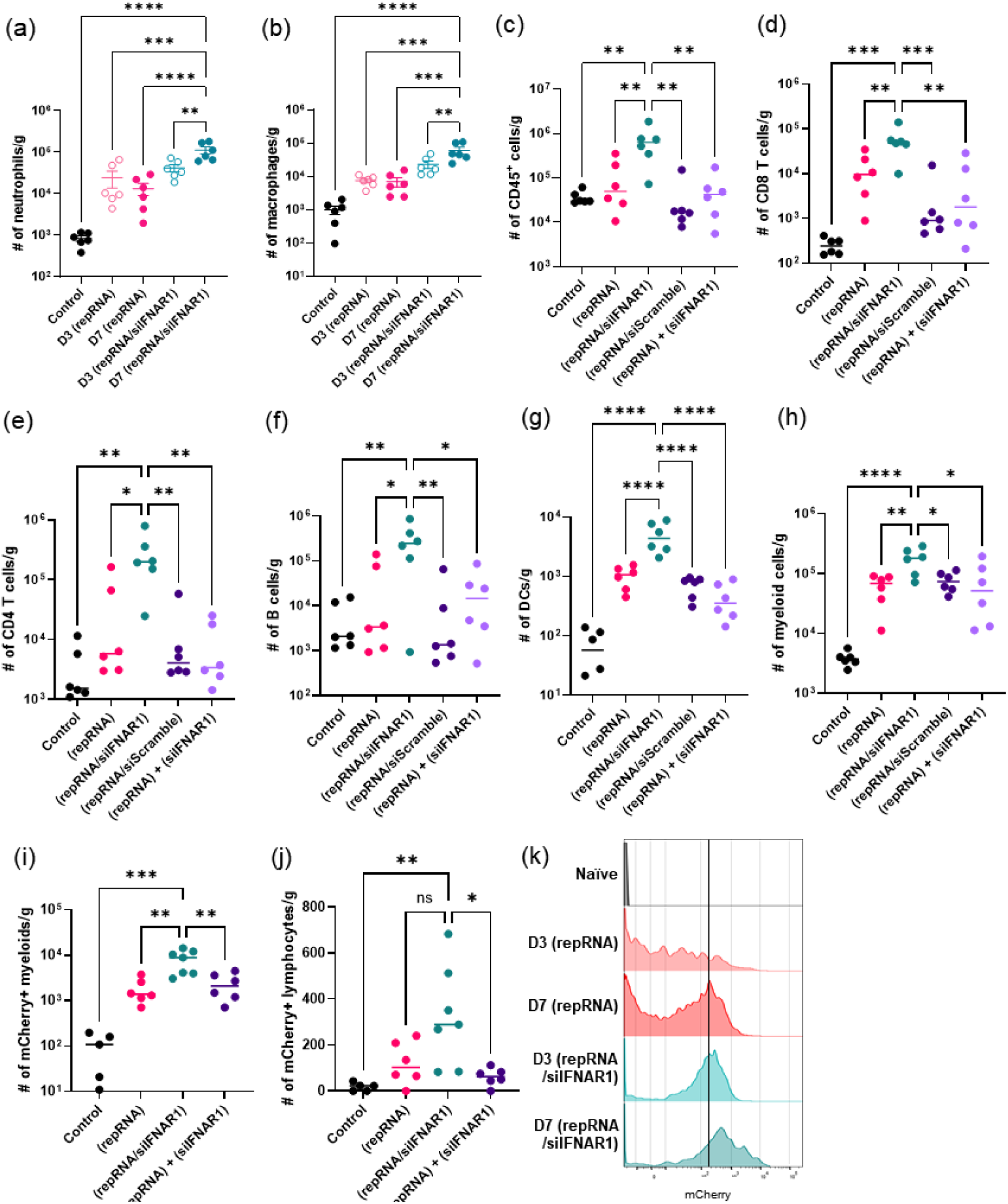
Ifnar1 silencing increases immune cell infiltration and activation in the muscle. Groups of Balb/C mice (n=5 animals/group) were immunized i.m. in each leg with 1 µg repRNA (encoding either immunogen or mCherry) loaded in LNPs, and sacrificed at days 3 or 7 post-injection for gastrocnemius muscle harvesting. Single cells were isolated from the muscles for flow cytometry-based immunophenotyping. (a-b) shows counts from the muscle of: (a) neutrophils; and (b) macrophages at days 3 and 7 post-vaccination by either repRNA alone or with siIFNAR1. (c-j) are immunophenotypes of mice at day 7 post-vaccination with repRNA alone, with siIFNAR1, with siScramble, or in cocktail mix with siIFNAR1-loaded LNPs. Shown are counts of: (c) CD45+ cells; (d) CD8+ T cells; (e) CD4+ T cells; (f) B220+CD19+ mature B cells; (g) MHCII+CD11c+ dendritic cells; (h) CD11b+ myeloid cells; (i) mCherry+ CD11b+ myeloids; and (j) mCherry+ lymphocytes. (k) shows MFI histogram of mCherry+ monocytes. Statistics represent one-way ANOVA and Tukey’s HSD Test (*, p<0.05; **, p<0.01; ***, p<0.001; ****, p<0.0001).

**Extended Data Figure 5.**
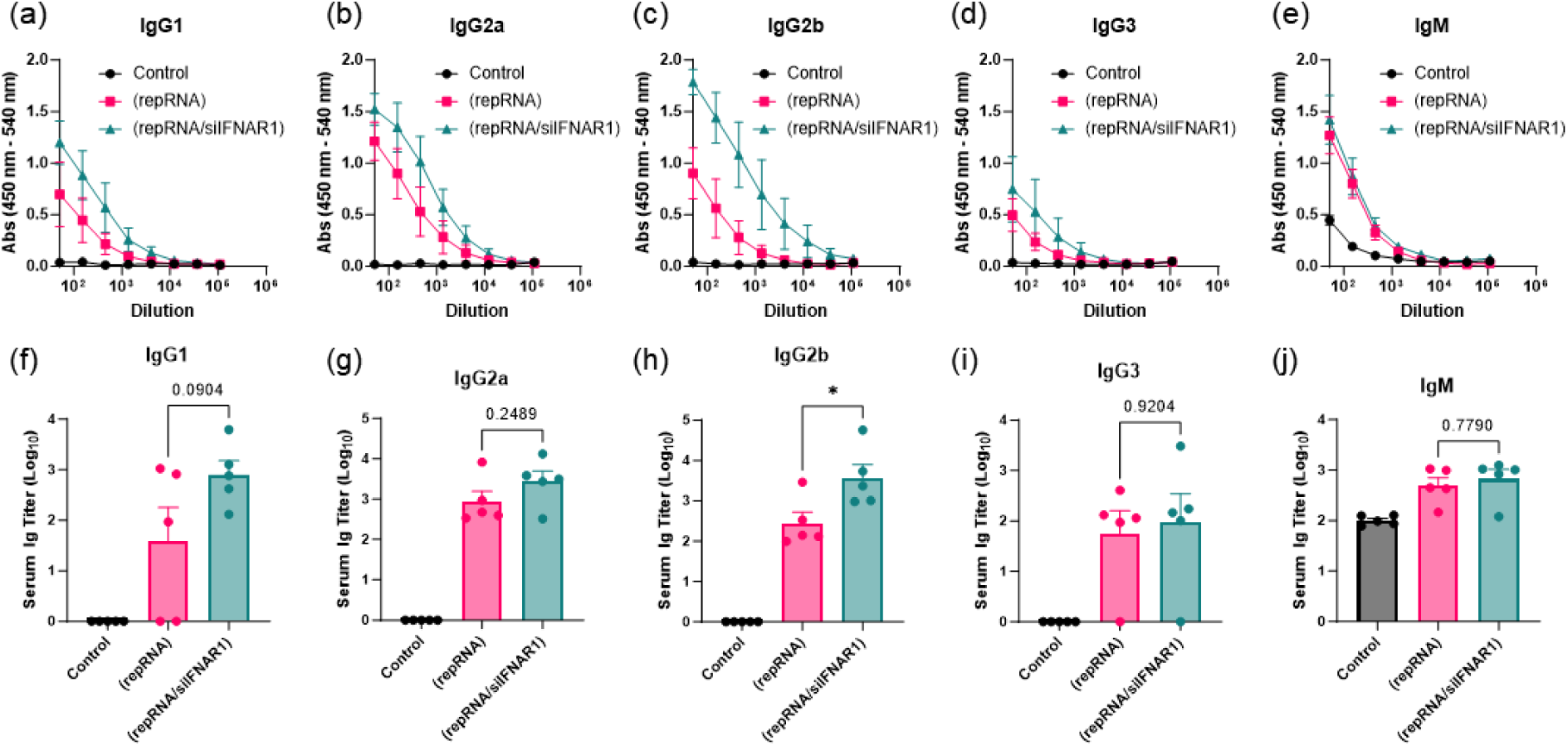
Antibody isotypes from mouse sera. Groups of Balb/C mice (n=5 animals/group) were immunized i.m. in each leg with 1 µg repRNA loaded in LNPs and were evaluated for vaccine-elicited immune responses. Shown are dilution curves (a-e) and endpoint serum Ig titers (f-j) at 2-weeks post-vaccination with specific isotypes: (a, f) IgG1; (b, g) IgG2a; (c, h) IgG2b; (d, i) IgG3; and (e, j) IgM. Statistics represent one-way ANOVA and Tukey’s HSD Test (*, *p*<0.05) of the endpoint serum Ig titers of each group.

**Extended Data Figure 6.**
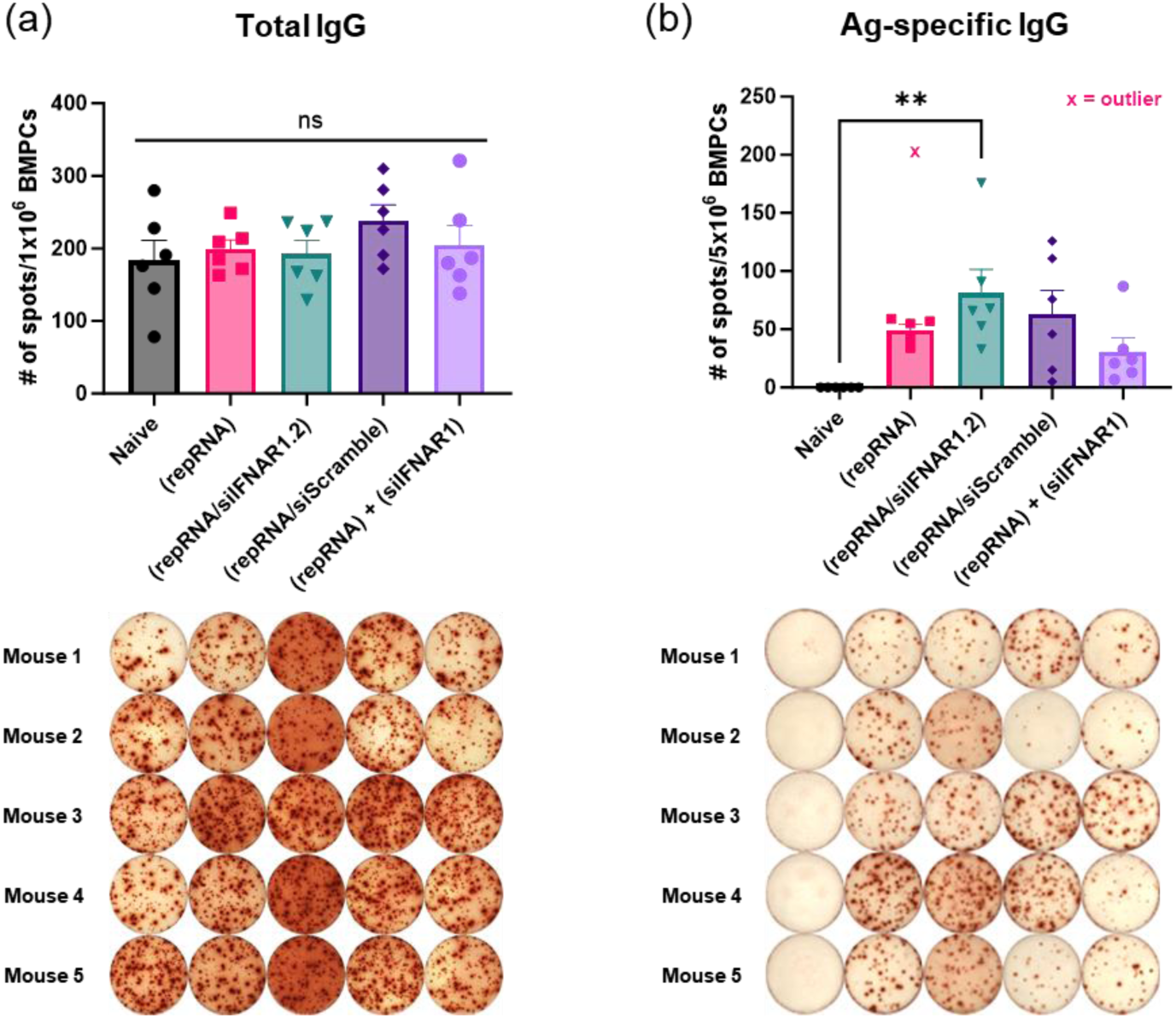
Changes in the long-lived plasma cells with respect to *Ifnar1* silencing. Groups of Balb/C mice (n=5 animals/group) were immunized i.m. in each leg with 1 µg repRNA loaded in LNPs and were evaluated for long-lived plasma cell responses at week 6 post-vaccination. Shown are spot counts (top) and well images (bottom) of total IgG (a) and antigen-specific IgG (b) secreted from bone-marrow plasma cells. Column graphs show means ± s.e.m. Statistics represent one-way ANOVA and Tukey’s HSD Test (ns= not significant; **, *p*<0.01). Outlier was statistically identified by ROUT test.

**Extended Data Figure 7.**
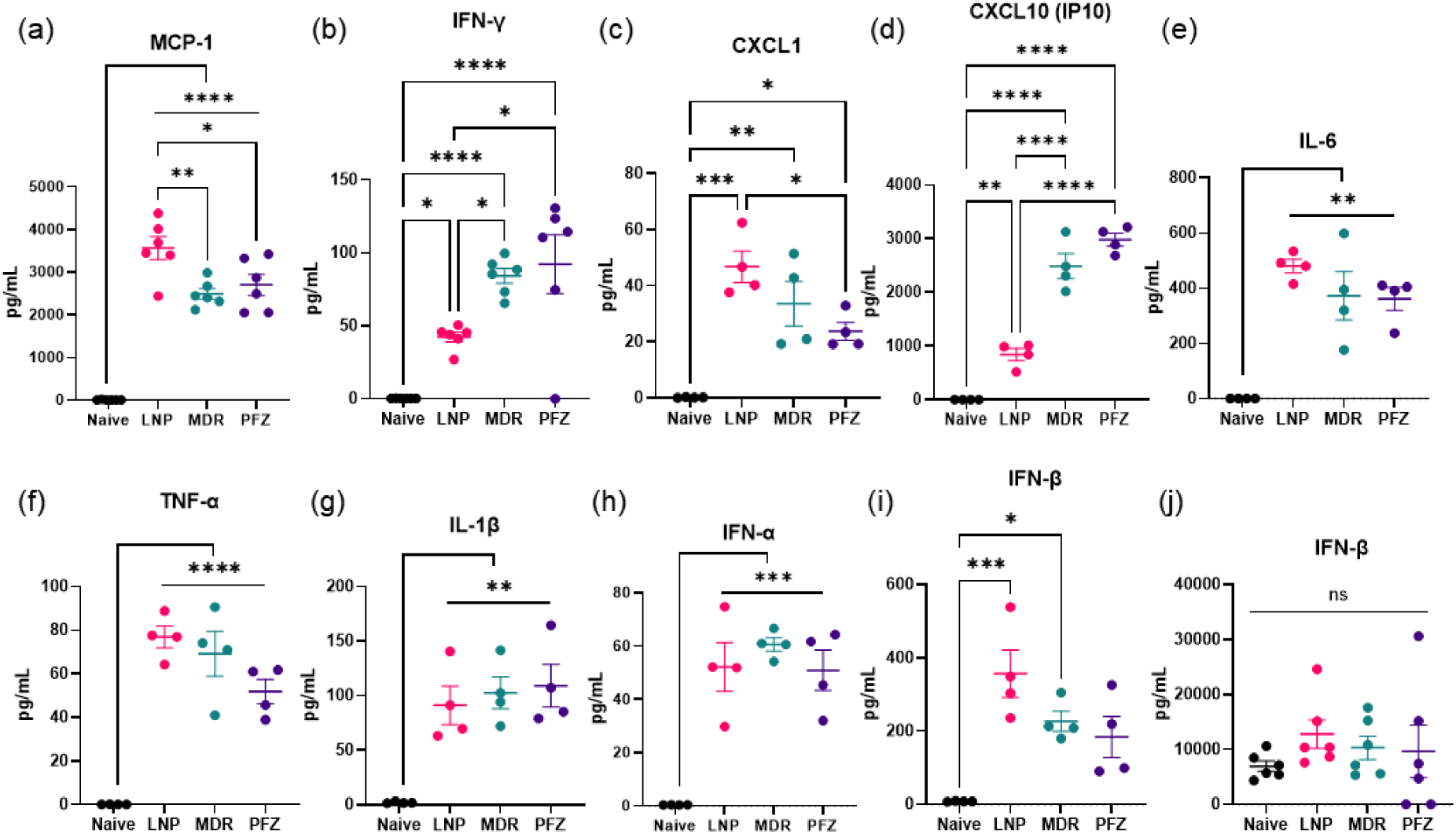
Cytokine profile in the muscle and draining lymph node from empty LNPs. Groups of Balb/C mice (n=5 animals/group) were immunized i.m. in each leg with empty LNPs and were evaluated for cytokine induction. Shown from the muscle are: (a) MCP-1; (b) IFN-γ; (c) CXCL1; (d) CXCL10; (e) IL-6; (f) TNF-α; (g) IL-1β; (h) IFN-α; and (i) IFN-β. From the popliteal lymph node, we show IFN-β (j) as a representative cytokine. Graphs show means ± s.e.m. Statistics represent one-way ANOVA and Tukey’s HSD Test (ns, not significant; *, p<0.05; **, p<0.01; ***, p<0.001; ****, p<0.0001).

**Extended Data Figure 8.**
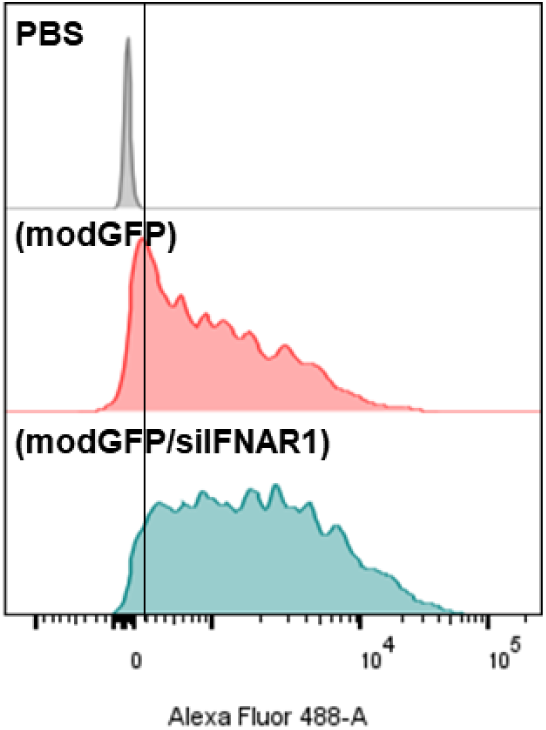
Effects of silencing *Ifnar1* on modRNA expression in C2C12 mouse myoblasts during *in vitro* transfection. C2C12 mouse myoblasts were treated with PBS, LNPs loaded with modRNA encoding for GFP (‘(modGFP)’), or LNPs that co-load modGFP and siRNA against *Ifnar1* (‘(modGFP/siIFNAR1)’). At 24h post-transfection, cells were analyzed by flow cytometry, shown as live cell histogram comparing expression of GFP.

**Extended Data Figure 9.**
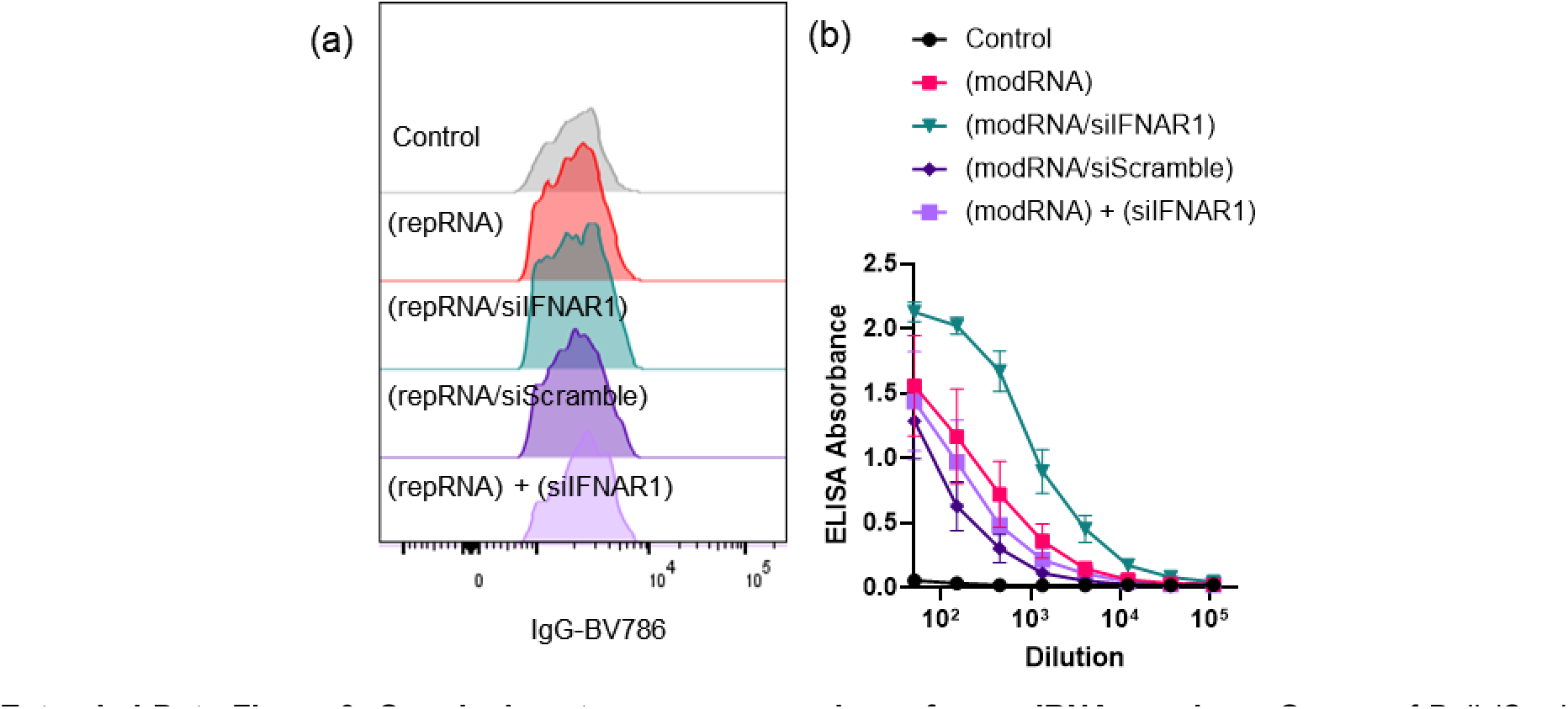
Germinal center response analyses for modRNA vaccines. Groups of Balb/C mice (n=5 animals/group) were immunized i.m. in each leg with 5 µg modRNA loaded in LNPs and were evaluated for vaccine-elicited immune responses. Shown are histogram of GC B cells expressing IgG at similar levels (a), and dilution curves of serum antibody at 2-weeks post-vaccination (b).

**Extended Data Figure 10.**
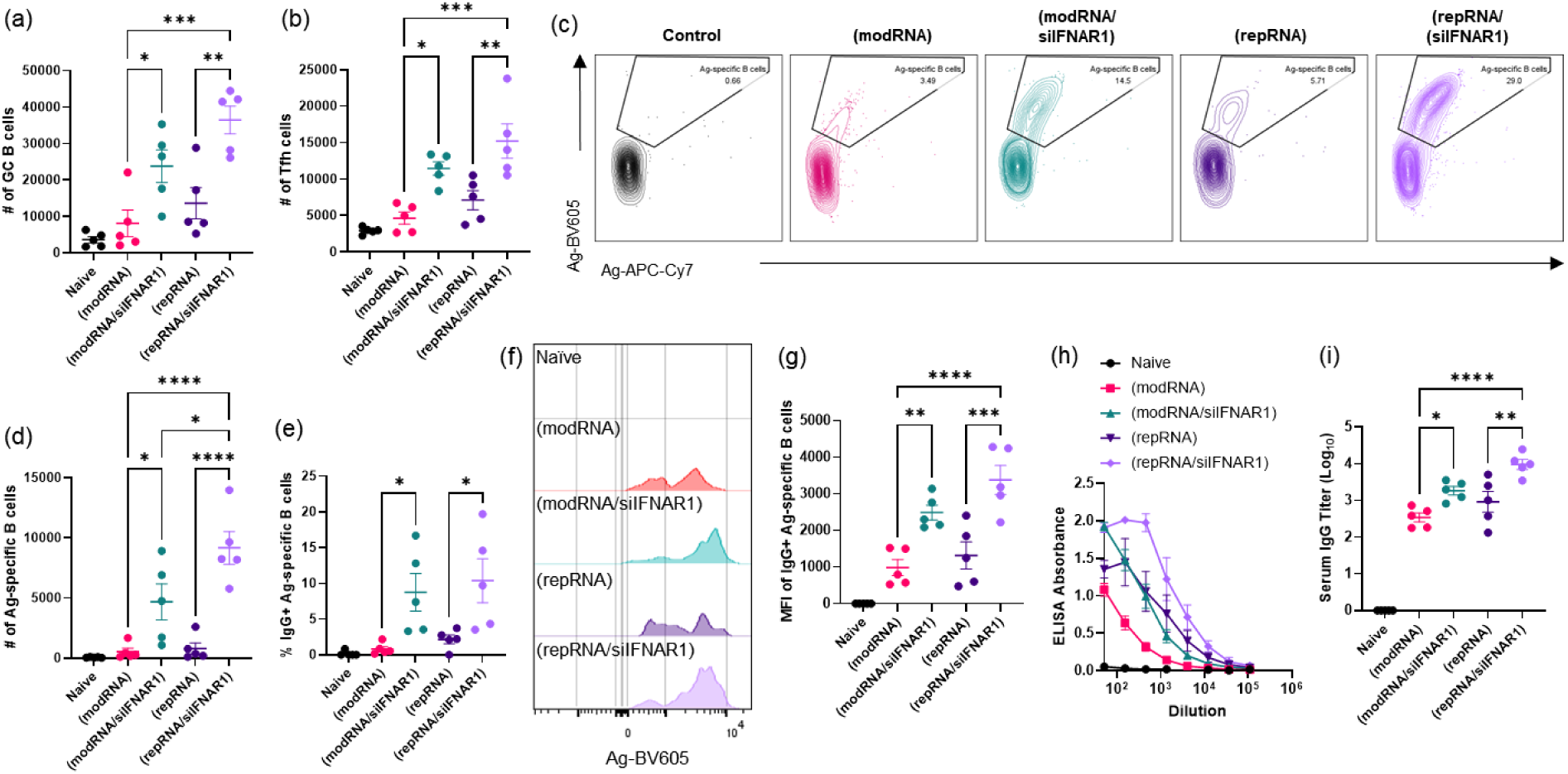
LNP formulation used in Moderna’s COVID-19 vaccine displays the same enhancements in vaccine-elicited immune response when co-delivering siIFNAR1. Groups of Balb/C mice (n=5 animals/group) were immunized i.m. in each leg with 1 µg repRNA or 5 µg modRNA loaded in the LNP formulation used in Moderna’s COVID-19 vaccine (mRNA-1273). Shown are counts of GC B cells (a), follicular helper T cells (b), gating for antigen-specific B cells (c), counts for antigen-specific B cells (d), and the frequency of IgG+ antigen-specific B cells (e). Histogram of antigen-specific B cells with equivalent IgG-expression levels is shown in (f), and the mean fluorescence intensity (MFI) of IgG+ antigen-specific B cells are shown in (g). Serum antibody responses were quantified by ELISA assay conducted on mouse sera collected at 14 days post-vaccination. The dilution curves are shown in (h), and the endpoint titer are shown in (i). Each column graph shows means ± s.e.m. Statistics represent one-way ANOVA and Tukey’s HSD Test (*, p<0.05; **, p<0.01; ***, p<0.001; ****, p<0.0001).

