## Supplementary Data for "Targeted suppression of type 1 interferon signaling during RNA delivery enhances vaccine-elicited immunity"

**Supplementary Table S1. Library of LNP formulations for intramuscular transfection**

| Label | Ionizable/Cationic (mol %) | | Helper (mol %) | | Cholesterol (mol %) | PEGylated lipid (mol %) | |
|---|---|---|---|---|---|---|---|
| LNP | TT3<br>Dlin-MC3-DMA | 10<br>25 | DOPE | 20 | 40 | DMG-PEG | 5 |
| MDR | SM-102 | 50 | DSPC | 10 | 38.5 | DMG-PEG | 1.5 |
| PFZ | ALC-0315 | 50 | DSPC | 10 | 38.5 | ALC-0159 | 1.5 |

| Supplementary Table S2. Physical properties of the synthesized vaccines | | | | | | | |
|---|---|---|---|---|---|---|---|
| **Payload** | **repRNA: siRNA (wt:wt)** | **RNA (ug)/dose** | **Lipid (ug)/dose** | **Encapsulation efficiency (%)** | **Mean size (nm)** | **PDI** | **ζ-potential (mV)** |
| repAg | 1:0 | 1 | 15 | 98.8 ± 2.0 | 44.3 ± 2.1 | 0.236 | 3.26 |
| repAg/siIFNAR1 | 1:2 | 3 | 15 | 97.4 ± 3.8 | 56.9 ± 1.8 | 0.215 | 0.87 |
| siIFNAR1 | 0:2 | 2 | 1.5 | 99.9 ± 0.1 | 47.1 ± 1.8 | 0.286 | -2.54 |

**Supplementary Table S3.** List of primers used for qRT-PCR

| Target | Forward | Reverse |
|---|---|---|
| repRNA Backbone | ATG CCG TAG GAC CAA ACT TC | TGG TGT CTA AAG CTG TCA GC |
| *Gfp* | AGT CCG CCC TGA GCA AAG A | TCC AGC AGG ACC ATG TGA TC |
| **Target** | **IDT DNA Assay ID** | **Ref Seq #** |
| *Gapdh* | Mm.PT.39a.1 | NM_008084(1) |
| *Ifnar1* | Mm.PT.58.33360098 | NM_010508(1) |

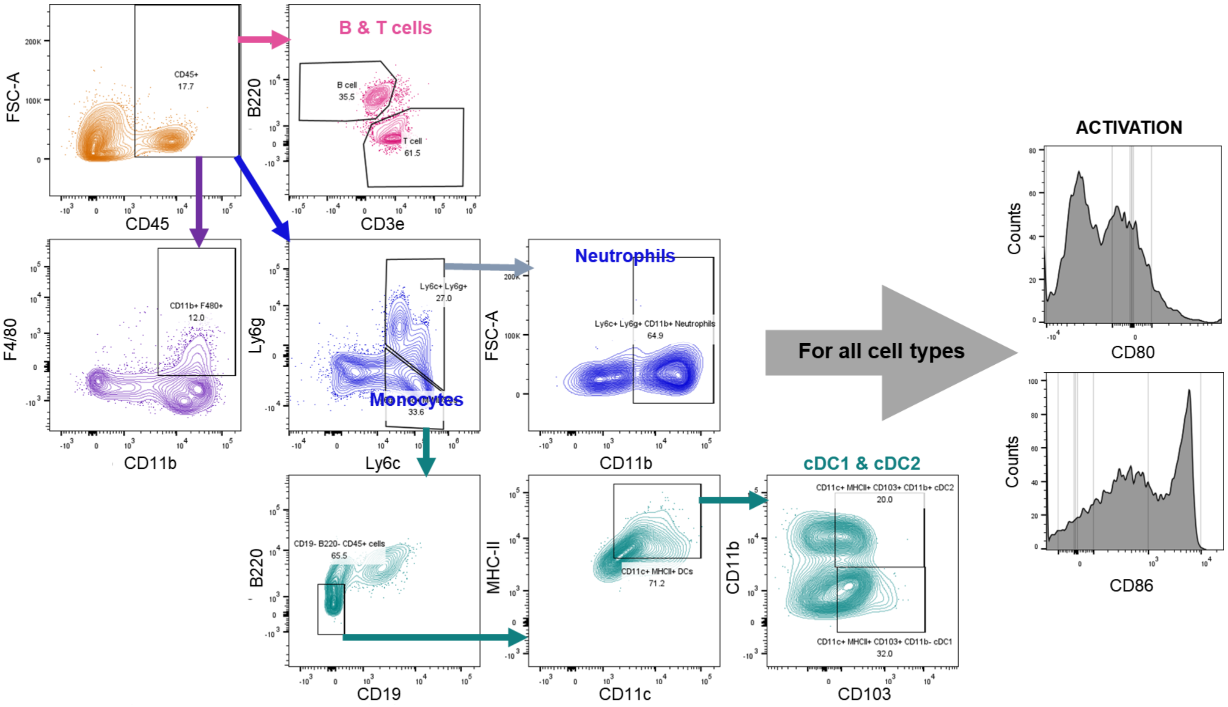


**Supplementary Figure S1. Gating strategy used for immunophenotyping.** Groups of balb/C mice (n=5 animals/group) were immunized i.m. in each leg with 1 µg repRNA (encoding either immunogen or mCherry) loaded in LNPs, and gastrocnemius muscle and popliteal lymph nodes were harvested. Single cells were analyzed for flow cytometry-based immunophenotyping of lymphocytes, myeloid cells and activation states.

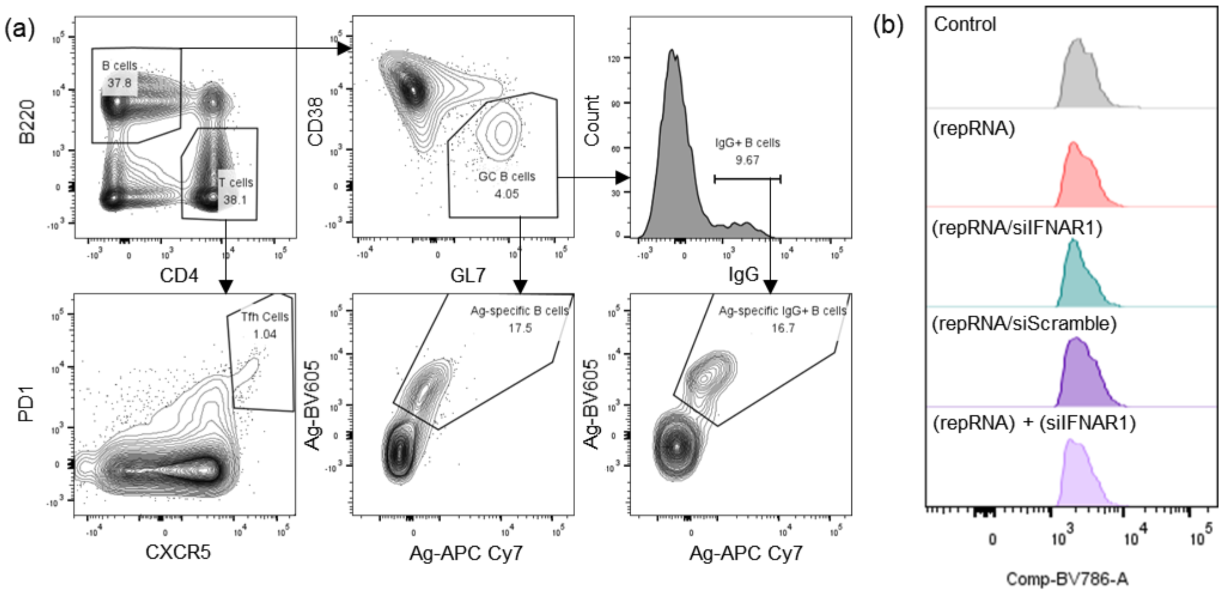


**Supplementary Figure S2. Germinal center response analyses.** Groups of balb/C mice (n=5 animals/group) were immunized i.m. in each leg with 1 µg repRNA loaded in LNPs and were evaluated for vaccine-elicited immune responses. Shown is the gating strategy (a), and histograms of GC B cells expressing IgG at similar levels (b).
